# Assembly of functionally antagonistic visual circuits for controlling pupil dynamics

**DOI:** 10.1101/500868

**Authors:** Onkar S. Dhande, Tania A. Seabrook, Ann H Phan, Lindsey D. Salaly, Nao Ishiko, Phong L. Nguyen, Jack T. Wang, Sylvia M. Evans, Andrew D. Huberman

## Abstract

Tbx20 plays a role in the development of non-image-forming pathways.
Loss of Tbx20-expressing RGCs alters specific aspects of the pupil reflex arc.
Previously unknown role for a specific subset of Tbx20-expressing RGCs in modulating pupil size.

**eTOC:** Dhande et al. identify a novel genetic program that marks and is required for the development of non-image-forming parallel visual pathways. Moreover, chemogenetic activation of a specific retina-to-OPN pathway reveals a novel circuit element controlling pupil size. These findings provide new insight into the mechanisms that establish defining features of functionally specialized sensory pathways.

**Summary:** The functional attributes of parallel sensory circuits are determined by the properties of the individual cell types that comprise them, and their connectivity. The mechanisms controlling the specification and establishment of parallel sensory pathways remain unclear. Here we report the non-uniform distribution of retinal ganglion cells (RGCs) expressing the transcription factor Tbx20 in mice. Owing to their layout within the retina, Tbx20-RGCs are positioned to preferentially encode light information from dorsal visual space. We discovered that Tbx20 expression is mostly restricted to three RGC-types: M1 intrinsically photosensitive RGCs (ipRGCs), M2 ipRGCs and ‘diving’ RGCs. The axonal projections of Tbx20-RGCs innervate non-image-forming brain centers including the suprachiasmatic nucleus, the medial division of the posterior pretectal nucleus, and the olivary pretectal nucleus (OPN), a principal station in the neural pathway for generating the pupillary light reflex (PLR). Conditional deletion of Tbx20 resulted in reduced Tbx20-RGC axonal innervation of these targets and revealed a key role of these neurons in driving specific phases of the PLR. Furthermore, chemogenetic activation uncovered a novel role for a specific subset of Tbx20-RGCs in controlling pupil dilation. These data offer a new understanding of the genetic and molecular mechanisms that establish specific, behaviorally-relevant visual circuits.

## Introduction

A hallmark feature of the mammalian eye-to-brain pathway is that the cells and circuits carrying specific aspects of visual information are often anatomically segregated from one another into parallel pathways. In the last decade, genetic tools have played a pivotal role in elucidating the identity of the specific retinal output neurons, the retinal ganglion cells (RGCs), and the targets they connect to in the brain (Hattar et al., 2002, Huberman et al., 2008; 2009; Kim et al., 2010; Kay et al., 2011; Dhande et al., 2013; 2015). What remains fundamentally lacking, however, is an understanding of how individual feature-detection channels within those parallel pathways are specified and established during development and how the RGCs within each discrete channel contribute to the operation of the circuit and consequently, how they drive behavioral outputs.

A prime example of a functionally dedicated pathway is the non-image-forming pathway serving the pupillary light reflex (PLR). The PLR allows the visual system to titrate the amount of light reaching the photoreceptors, reduce optical aberrations, and increase the effective focus, thereby optimizing visual sensitivity and acuity (Denton 1956; Campbell & Gregory, 1960). The primary stages of the PLR involve eye-to-brain signaling from a dedicated set of RGCs that relay changes in overall ambient light levels reaching the eye, to neurons in the OPN - a bilaterally symmetric set of targets in the dorsal midbrain (Trejo & Cicerone 1984; Young and Lund 1994; Gamlin and Clarke 1995; Gamlin et al. 1995; Distler & Hoffmann 1989a, b).

A specific category of retinal projection neurons termed ‘intrinsically photosensitive RGCs’ (ipRGCs) is crucial for the integrity of the PLR. The ipRGCs are so-named because they not only encode light signals from rod/cone photoreceptors but also express the photopigment melanopsin/OPN4 and thus are endowed with the property of responding directly to light (i.e. melanopsin-dependent depolarization). As a group, ipRGCs include five subtypes, ‘M1-M5’, each with different morphologies, degrees of photosensitivity, patterns of central connections and thus functional roles (Berson et al., 2002; Hattar et al., 2002; Sand et al., 2012). Previous work demonstrated that the axons from different ipRGC subtypes are compartmentalized within the OPN. The M1 ipRGCs express the highest levels of melanopsin and thus display the strongest intrinsic photosensitivity and project their axons to the ‘shell’ of the OPN, whereas the M2 ipRGCs and the ‘diving’ RGCs project to the OPN ‘core’ (Fig. 1A-A”, Schematized in Fig. 1B) (Hattar et al., 2002, 2006; Baver et al., 2008; Osterhout et al., 2011; 2014). The shell-projecting M1 ipRGCs are necessary for generating a normal light-driven PLR in response to changes in environmental light levels; ablating these M1s drastically attenuates the PLR (Güler et al., 2008). Given that M1s project to the OPN shell, this raises the question: what is the functional relevance of retino-OPN core connectivity? Indeed, the functional role of the OPN core remains unknown and the specific output channels of the shell versus core remain mysterious, as does their greater role in driving the PLR and other behaviors.

**Figure 1:**
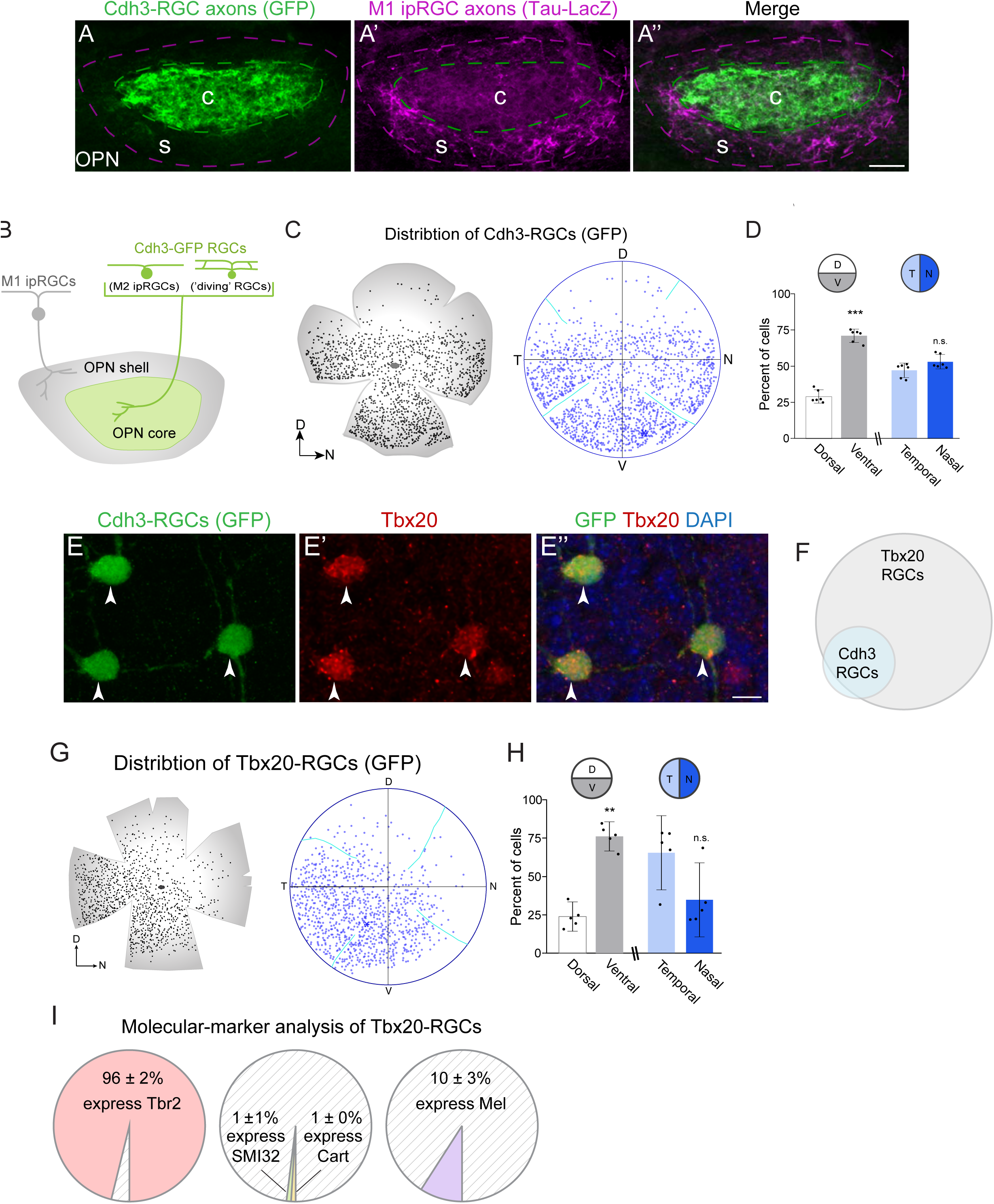
Cdh3-RGCs form a unique non-image-forming optic pathway. (A-A’’) Photomicrographs showing Cdh3-RGC projections (green, GFP) to the OPN core (c), and M1 ipRGC projections (purple, β-Gal immunostaining) to the OPN shell (s). (B) Schematic of the diversity of RGC types innervating the OPN. Cdh3-RGCs include two RGCs subtypes: M2 ipRGCs and ‘diving’ RGCs. (C) Retinal flat mount showing the ventral-biased spatial distribution of Cdh3-RGCs (GFP, black dots, left). Azimuthal equilateral projection of reconstructed retina with labeled Cdh3 cells (blue circles, right). Blue asterisk is the peak density. Cyan lines indicate stitching locations from flat mount relieving cuts. (D) Percent of Cdh3-RGCs (GFP) cells in respective retinal halves: dorsal (29.0 ± 1.8), ventral (71.0 ± 1.8), temporal (47.0 ± 2.0) and nasal (53.0 ± 2.0) (Cdh3-GFP RGCs/retina: 1005 ± 74; n= 6 retinas/mice, P20). *** p<0.001; n.s., not significant, p>0.05. Two-tailed paired Student’s t-test. Data shown are Mean ± SEM. (E-E’’) Photomicrograph of Cdh3-RGCs (green, E) co-immunostained with Tbx20 (red, E’) at P4. Cell nuclei are labeled with DAPI (blue, E’’). Arrowheads indicate overlap of GFP and Tbx20. (F) Venn diagram showing overlap of Cdh3-RGCs and Tbx20-RGCs. (83 ± 3% of Cdh3-RGCs express Tbx20, 562 GFP (Cdh3-RGCs) cells counted; 38 ± 1% of Tbx20 expressing cells are Cdh3-RGCs, 1242 Tbx20-expressing cells counted; data are Mean ± SEM; n=2 mice/retinas) (G) Retinal flat mount showing ventral-biased spatial distribution of Tbx20-RGCs (GFP, black dots, left). Azimuthal equilateral projection of reconstructed retina with labeled Tbx20 cells (blue circles, right). Cyan lines indicate stitching locations from flat mount relieving cuts. (H) Percent of Tbx20-GFP cells in respective retinal halves: dorsal (23.9 ± 3.1), ventral (76.1 ± 3.1), temporal (65.4 ± 7.9) and nasal (34.6 ± 7.9) (Tbx20-RGCs/retina: 863 ± 54; n=5 retinas/mice, P20). *** p<0.01; n.s., not significant, p>0.05. Two-tailed paired Student’s t-test. Data shown are Mean ± SEM. (I) Quantification of expression of RGC type-specific markers by Tbx20-RGCs. Data shown are Mean ± SEM. (Tbr2: n=3 mice/retinas, 374 Tbx20-GFP cells counted; Cart: n=2 mice/retinas, 282 Tbx20-GFP cells counted; SMI32: n=2 mice/retinas, 184 Tbx20-GFP cells; Melanopsin: n=4 mice/retinas, 693 Tbx20-GFP cells) See also Figure S1 and Figure S2. D: dorsal; N: nasal; V: ventral; T: temporal. Scale bar: 100μm (A’’), 10μm (E’’)

Here we used molecular genetic tools centered around the T-box transcription factor Tbx20 to visualize a specific subset of non-image-forming RGCs including those that project from the eye to the ‘core’ compartment of the OPN, along with cell-type-specific profiling, conditional knockout of Tbx20 and *in vivo* behavioral analyses, to parse the organizational logic and mechanisms underlying the specification of non-image forming visual circuits. We discovered a molecularly defined subset of RGCs that are arranged in a unique distribution across the retinal surface and project to distinct target regions within the non-image-forming pathway. We then used conditional alleles of Tbx20 and chemogenetic activation to discover that Tbx20 plays a crucial role in establishing specific non-image-forming pathways which we find modulate pupil size. These findings are discussed in terms of the potential role of this circuit in visual behaviors and in the context of understanding how parallel sensory pathways are designated and assembled.

## Results

### Cdh3-RGCs are non-uniformly distributed in the retina

In the transgenic mouse line Cdh3-GFP, the retina expresses enhanced GFP (GFP) in a subset of M2-type ipRGCs and ‘diving’ RGCs (hereafter referred to as Cdh3-RGCs), which project their axons to a specific set of non-image-forming centers in the brain: the OPN-core, medial division of the posterior pretectal nucleus (mdPPN) and ventral lateral geniculate nucleus (vLGN) (Osterhout et al., 2011; 2014). The spatial distribution of specific RGC-types across the retina has direct implications for the kind of visual information processed by their respective target in the brain. Therefore, we systematically examined the distribution of Cdh3-RGCs (GFP^+^) and observed they are arranged in a striking ventral:high, dorsal:low density-distribution across the retinal surface (Fig. 1C). The distribution of Cdh3-RGCs was significantly biased towards the ventral half of the retina (p<0.001, Fig. 1D left). However, no significant bias was observed for the distribution of Cdh3-RGCs between the nasal and temporal half of the retina (p=0.2; Fig. 1D right). This ventrally-biased layout indicates that Cdh3-RGCs are poised to encode visual features preferentially from the dorsal visual field. Furthermore, this also implies that target nuclei receiving input from these RGCs are positioned to extract visual information from the dorsal visual field. Together, the characteristics of Cdh3-RGCs and their central projections to the midbrain reveal a unique subdivision of the retinofugal pathway.

### Tbx20, a T-box family transcription factor, is highly enriched in Cdh3-RGCs

The identification of molecular markers for M1 ipRGCs has greatly facilitated the study and understanding of their development and function (McNeill et al., 2011; Chen et al., 2011; Fox & Guido, 2011). Similarly, to further understanding of the retino-OPN pathway we performed an immunohistochemical screen to identify molecular markers for Cdh3-RGCs, focusing specifically on transcription factors. Those experiments revealed Tbx20, a T-box family member, as an intriguing candidate marker for Cdh3-RGCs.

To determine the expression of Tbx20 in Cdh3-RGCs, we stained whole mount retinas from Cdh3-GFP mice with an antibody specific to Tbx20 protein (Song et al., 2006). More than 80% of Cdh3-RGCs expressed Tbx20 at postnatal day 4 (P4) (Fig. 1E-E”, 1F). Conversely, Cdh3-RGCs comprised a smaller fraction (∼38%) of the Tbx20 expressing population of retinal neurons (Fig. 1F). To determine the other types of RGCs that express Tbx20 and their distributions across the retina, we also used a GFP mouse line expressing a Tbx20-GFP fusion protein, which recapitulates the endogenous nuclear expression of Tbx20 (Shen et al., 2011). First, examination of the spatial arrangement of Tbx20-RGCs (Tbx20-GFP-expressing) revealed a ventral: high, dorsal: low gradient of Tbx20-RGC soma density (Fig. 1G, H, and Fig. S1). The biased distribution of Tbx20-RGCs was already apparent in flat-mounted retinas on embryonic day (E) 18.5 (Fig. S1A-A2), a developmental time point when most RGCs have differentiated (Young 1985) and persisted through the early postnatal weeks and into adulthood (Fig. S1B-D). We found the distribution of Tbx20-RGCs at P20 was significantly biased towards the ventral half of the retina, similar to Cdh3-RGCs (p=0.0016, Fig. 1H left). Although there was a strong temporal bias in the distribution of Tbx20-RGCs in 4/5 retinas, on average the distribution of Tbx20-RGs in temporal versus nasal retina was not significantly different (p=0.2, Fig. 1H right). Thus, both Tbx20-RGCs and Cdh3-RGCs have a similar dorsal:low to ventral:high gradient, although Tbx20-RGCs have a qualitative bias towards the ventral-temporal quadrant (compare Fig. 1G, H to Fig. 1C, D). These data suggest that the non-uniform distribution of Tbx20-RGCs is an early and persistent feature of this cell type, and not the transient consequence of postnatal events that shape retinal circuitry (e.g., naturally occurring developmental cell-death) (Young 1984; Crespo et al., 1985; Williams et al., 1986).

Second, by staining retinas from Tbx20-GFP mice for other markers of RGC identity, we found that almost all Tbx20-RGCs express Tbr2 (∼96%; Fig. S2A-A”, 2F), which is a marker of non-image-forming RGCs (Mao et al., 2014; Sweeney et al., 2014, 2017; Seabrook et al., 2017). Additionally, we found that many Tbx20-RGCs express Brn3b (Mean ± SEM: 76 ± 6%, n=4 mice/retinas, 465 Tbx20-GFP cells; data not shown), a transcription factor known to play a role in the development of non-image-forming RGCs, including ipRGCs, and their central projections (Quina et al. 2005; Badea et al., 2009). A negligible number of Tbx20-RGCs express Cart (∼1%), a marker of On-Off direction-selective RGCs (Kay et al., 2011), or SMI32 (∼1%), a marker of alpha-like RGCs (Lim et al., 2007) (Fig. S2B-C’’, 2F). A small fraction of Tbx20-RGCs (∼10%) express the photopigment melanopsin at high enough levels to be detected by standard immunohistochemistry (Fig. S2D-D”, 2F). Together, these data suggest that Tbx20 is expressed by a mixed but relatively restricted subset of non-image-forming RGCs, including those projecting to the OPN core.

### Tbx20 loss results in reduced retinal innervation of non-image-forming targets

Does Tbx20 play a role in establishing non-image-forming pathways? To address this, we used the Cre-loxP system to genetically delete Tbx20 from non-image-forming RGCs. We screened the GENSAT repository of BAC transgenic mice (Gong et al., 2003) to identify lines in which Cre recombinase was expressed in non-image-forming RGCs. From that screen, we identified the Tph2-Cre mouse line (Cre^+^) as a potential candidate. We found that ∼95% of Tbx20-RGCs express Cre recombinase in the Tph2-Cre mouse line (Fig. 2A-B). Moreover, we previously reported that in the Tph2-Cre mouse line the majority of non-image-forming RGCs express Cre [Fig. 2C-C”; previously quantified: 84% of melanopsin^+^ ipRGCs, 82% of Tbr2^+^ RGCs, and 90% of Cdh3-RGCs express Tph2-Cre (Seabrook et al., 2017)]. Thus, we crossed Cre^+^ mice to transgenic mice harboring floxed alleles of Tbx20 (Tbx20^f/f^) in order to generate heterozygous Tbx20 conditional knockout (cKO) offspring (Cre^+^; Tbx20^f/+^). These mice were in turn crossed to Tbx20^f/f^ to produce Tbx20 conditional knockout mice (Cre^+^; Tbx20^f/f^) (hereafter referred to as ‘Tbx20 cKO mice’ for simplicity).

**Figure 2:**
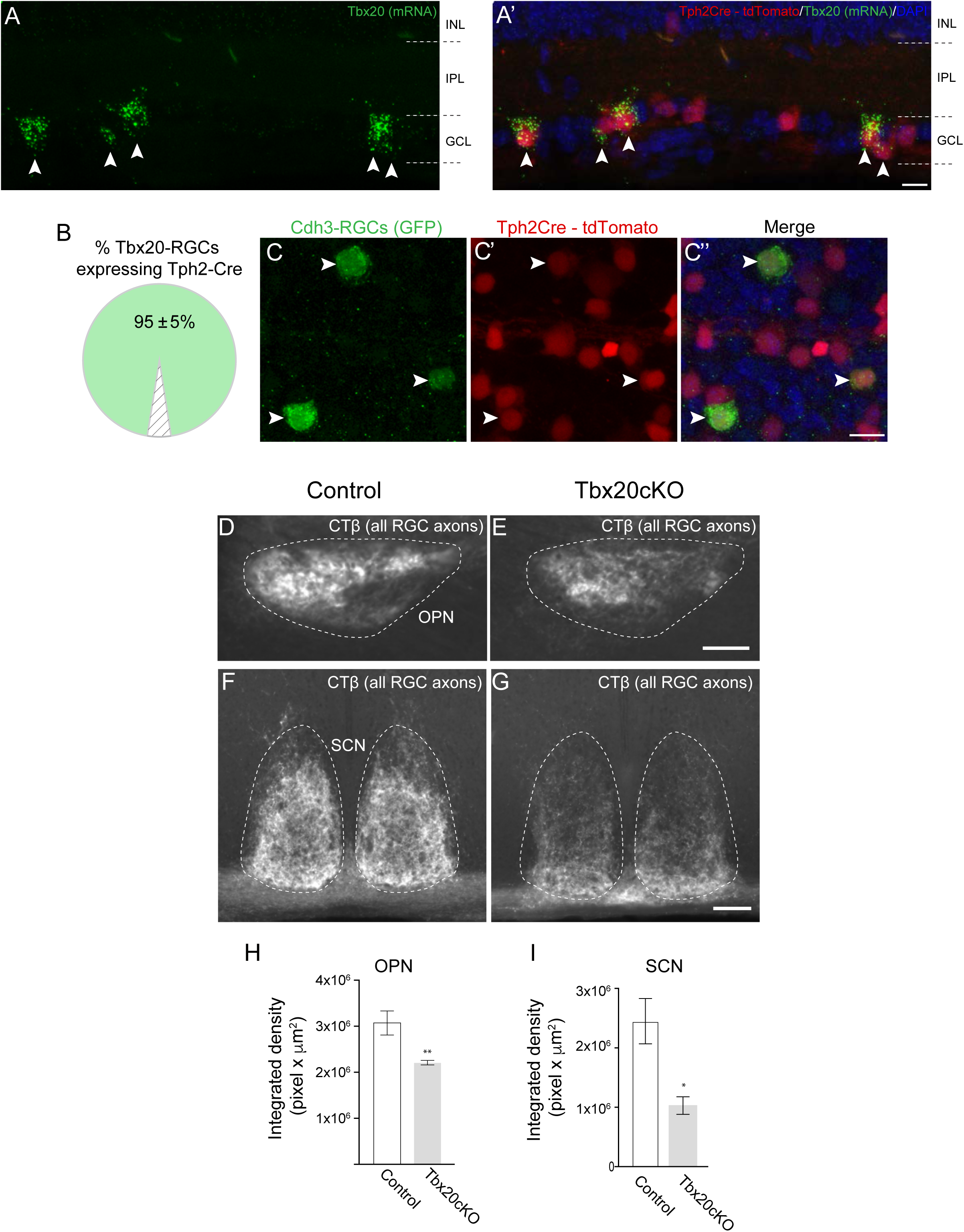
Loss of Tbx20 results in reduced retinal innervation of non-image-forming targets. (A-A’) Photomicrographs showing cells expressingTbx20 mRNA (green, A) also express Cre recombinase (visualized by tdTomato reporter, endogenous signal, red A’) at P19. Nuclei are labeled with DAPI (blue, A’). Arrowheads indicate overlap of Tbx20 mRNA and tdTomato (Cre). (B) Quantification of expression of Cre recombinase (tdTomato reporter) by Tbx20-RGCs. Data shown is Mean ± SEM. (n=2 mice/retinas, 153 Tbx20 cells) (C-C’’) Photomicrographs showing Cdh3-RGCs (green, C) overlap with Cre recombinase (visualized by tdTomato reporter, red, C’) expressing cells. Nuclei are labeled with DAPI (blue, C’’) Arrowheads indicate overlap of GFP and tdTomato (Cre). (D-G) Example photomicrographs showing RGC innervation (visualized by CTβ labeling) of OPN and SCN in control and Tbx20 cKO mice. (H, I) Quantification of density of retinofugal innervation in OPN (D, E) (Control Mean ± SEM: 3.1 x10^6^ ± 2.6 x10^5^ pixel x µm^2^, n= 6 mice; Tbx20 cKO Mean ± SEM: 2.2 ×10^6^ ± 4.8×10^4^ pixel x µm^2^, n=6 mice) and SCN (F, G) (Control Mean ± SEM: 2.5 ×10^6^ ± 3.8 ×10^5^ pixel x µm^2^, n= 6 mice; Tbx20 cKO Mean ± SEM: 1.0 x10^6^ ± 1.5×10^5^ pixel x µm^2^, n=4 mice) in control and Tbx20 cKO mice at P20. See also Figure S3 and Figure S4. GCL: ganglion cell layer; IPL: inner plexiform layer; INL: inner nuclear layer; OPN: olivary pretectal nucleus; SCN: suprachiasmatic nucleus. Scale bar: 10μm (A”, C”), 100μm (E, G). * p <0.05, ** p<0.01. Student’s t-test.

To assess the consequence of deleting Tbx20 on the non-image-forming pathways we first visualized the entire retinofugal pathway in control and Tbx20 cKO mice by injecting the anterograde fluorescent tracer CTβ-488 into both eyes to label RGC axons. The OPN, a major target of Cdh3-RGCs and M1 ipRGCs, and the suprachiasmatic nucleus (SCN), a major target of M1 ipRGCs, were clearly identifiable based on CTβ^+^ RGC axonal labeling in Tbx20 cKO mice (Fig. 2D-G). The OPN and SCN area occupied by CTβ^+^ RGC axons was not significantly different between control and Tbx20 cKO mice (data not shown). However, the innervation density of RGC axons in the OPN (core and shell) and the SCN was significantly reduced in Tbx20 cKO mice, in comparison to control mice (OPN: ∼28% reduction, p=0.008; SCN: ∼ 58% reduction, p=0.019; Fig. 2H, I). These data suggest that the loss of Tbx20 function leads to a disruption of the parallel pathways forming the non-image-forming system.

### A role for Tbx20 in establishing the retino-OPN-core parallel pathway

What is the consequence of deleting Tbx20 on OPN-core-projecting RGCs? To visualize this we generated Tbx20 cKO mice on a Cdh3-GFP background, yielding Tph2-Cre^+^; Cdh3-GFP^+^; Tbx20^f/f^ triple transgenic/conditional mutant mice. By examining the axonal projections of Cdh3-RGCs in the brains of juvenile Tbx20 cKO mice and in littermate-control mice (Tph2-Cre^+^; Cdh3-GFP^+^; Tbx20^f/+^, and Cdh3-GFP^+^; Tbx20^f/f^, and Cdh3-GFP^+^; Tbx20^f/+^; P19-P25) we found that, in the absence of Tbx20 function, Cdh3-RGC innervation of the OPN-core was substantially reduced. In 2/7 Tbx20 cKO mice, the projections were modestly reduced (Fig. 3Q, 3R) and in 5/7 mice, the projections to this target were completely absent (Fig. 3B-E, 4P, schematized in Fig. 3Y). The same general effect was observed for the mdPPN, a target situated immediately caudal to the OPN. Normally, the mdPPN is densely innervated by Cdh3-RGC axons (Fig. 3F), but after conditional removal of Tbx20 from Cdh3-RGCs, the mdPPN was also largely devoid of Cdh3-RGC axonal innervation (Fig. 3G-J, 3S-U, schematized in Fig. 3Y). Cdh3-RGC inputs to the vLGN-another brain target that normally receives substantial input from Cdh3RGCs, were absent or substantially reduced in Tbx20 cKO mice (Fig. 3L-O, 3V-X).

**Figure 3:**
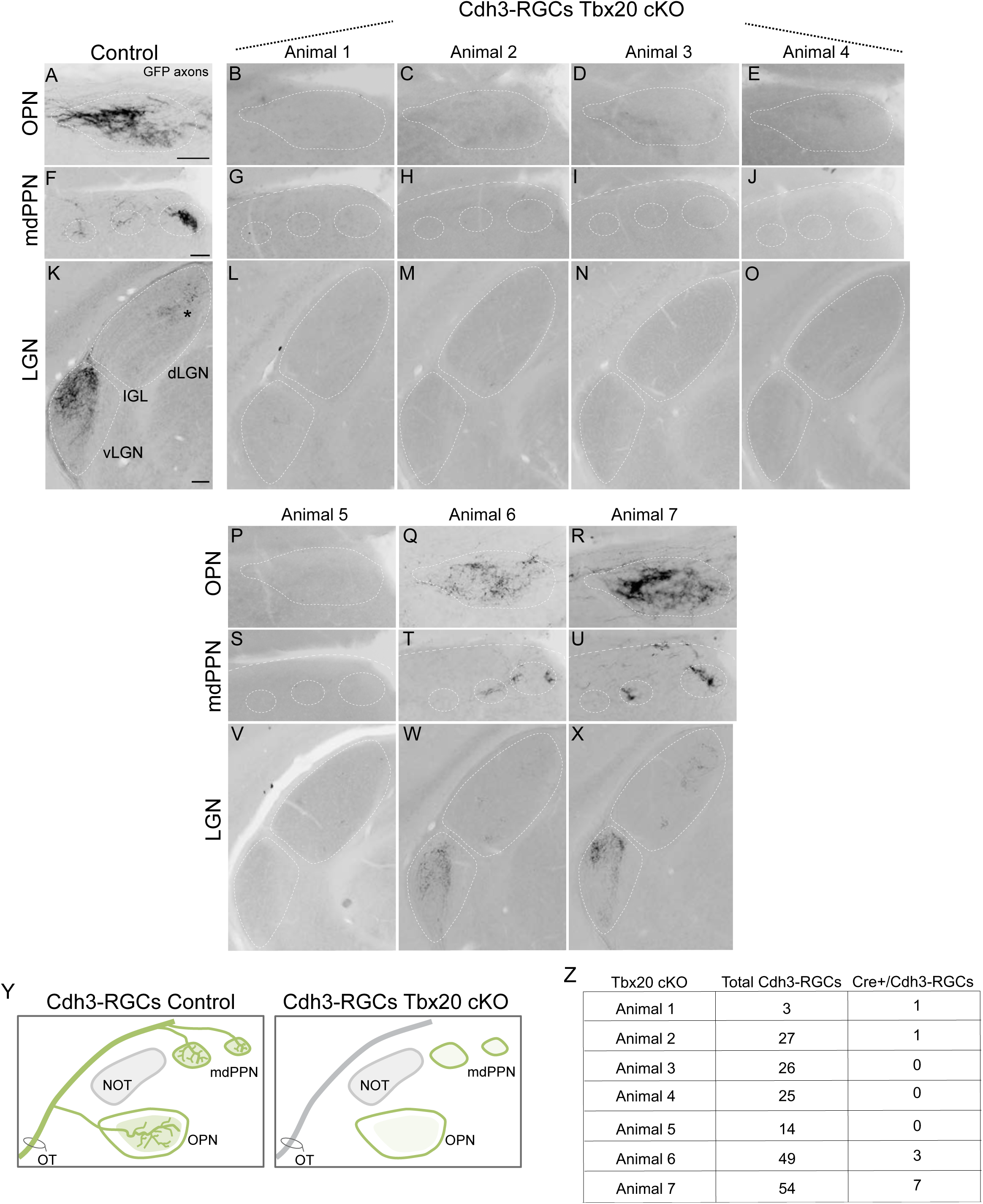
Conditional deletion of Tbx20, a transcription factor highly enriched in Cdh3-RGCs, results in loss of Cdh3-RGC projection pathway. (A, F, K) Photomicrographs showing innervation of OPN and mdPPN by Cdh3-RGC axons (black) in control animals. In the visual thalamus, Cdh3-RGC axons selectively terminate in the vLGN and sparsely innervate the dorsal-medial half of the dLGN (asterisk). (B-E, G-J, L-O, P-X) Photomicrographs demonstrating the variability in the lack of innervation of OPN, mdPPN and LGN by Cdh3-RGC axons in Tbx20 cKO mice. (Y) Schematics depicting typical outcome of Cdh3-RGC innervation of their normal targets in control and Tbx20 cKO mice. (Z) Quantification of total number of Cdh3-RGCs and fraction of Cre expressing Cdh3-RGCs remaining in Tbx20 cKO mice. The level of OPN, mdPPN and LGN innervation by Cdh3-RGC axons in Tbx20 cKO mice is strongly correlated with the number of spared (Cre negative) Cdh3-RGCs. OPN: olivary pretectal nucleus; mdPPN: medial division of the posterior pretectal nucleus; dLGN: dorsal lateral geniculate nucleus; vLGN: ventral LGN; IGL: intergeniculate leaflet. Scale bar: 100μm (A, F, K).

Is the lack of target innervation by Cdh3-RGCs observed in Tbx20 cKO mice due to a loss of Cdh3-RGCs? To address this, we first quantified the number of Cdh3-RGCs present in Tbx20 cKO retinas (see methods). We found that only a small number of Cdh3-RGCs remained in Tbx20 cKO mice (Mean ± SEM: 30 ± 8, n=7 mice/retinas; Fig. 3Z). This indicates that conditional deletion of Tbx20 results in a severe reduction in Cdh3-RGCs. Moreover, >90% of the spared Cdh3-RGCs in Tbx20 cKO retinas did not express Cre [confirmed by the absence of tdTomato (Cre-dependent reporter); Fig. 3Z] and thus, any Cdh3-RGCs spared was the consequence of lack of Cre dependent Tbx20 excision, not a failure for Tbx20 deletion to eliminate these cells. The absence of Cdh3-RGC cells in Tbx20 cKO retinas suggests that the reduction in Cdh3-RGC inputs to the OPN-core observed in Tbx20 cKO mice reflects the loss of Cdh3-RGCs altogether. However, these data cannot exclude the possibility that loss of Tbx20 might somehow result in a loss of GFP expression in Tbx20 cKO mice.

### Loss of Tbx20 partially disrupts the organization of the retino-OPN-shell pathway

It is well established that M1 ipRGCs express the highest levels of the photopigment melanopsin relative to all other ipRGC types and as a result, display the greatest intrinsic light responses (Schmidt et al., 2011; Sand et al., 2012). Furthermore, M1 ipRGCs are the exclusive source of retinal inputs to the OPN shell-the pathway thought to drive the PLR (Hattar et al., 2002; Viney et al., 2007; Baver et al., 2008; Berson et al., 2010; Güler et al., 2008). A small fraction of Tbx20-RGCs express high levels of melanopsin (Fig. 1I, Fig. S2D-D’); therefore it seemed reasonable to assume that some of those Tbx20-RGCs might be M1 ipRGCs. Hence, we sought to determine if Tbx20 function is necessary for normal development of the retino-OPN-shell pathway.

As a first step toward that goal, we stained flat-mounted retinas from Tbx20 cKO mice and littermate-control mice with an antibody against melanopsin to visualize M1 ipRGCs. The somas and dendrites of M1 ipRGCs show the strongest immunoreactivity for melanopsin (Berson et al., 2010; see Methods for details on antibody specificity). In Tbx20 cKO mice, the layout of the network of ipRGC types was severely disrupted, although some M1 ipRGCs were still present (Fig. 4A-B’, schematized in Fig. 4C). The severe but incomplete loss of M1 ipRGCs is consistent with our finding that not all ipRGCs that express high levels of melanopsin also express Tbx20 protein (see above).

**Figure 4:**
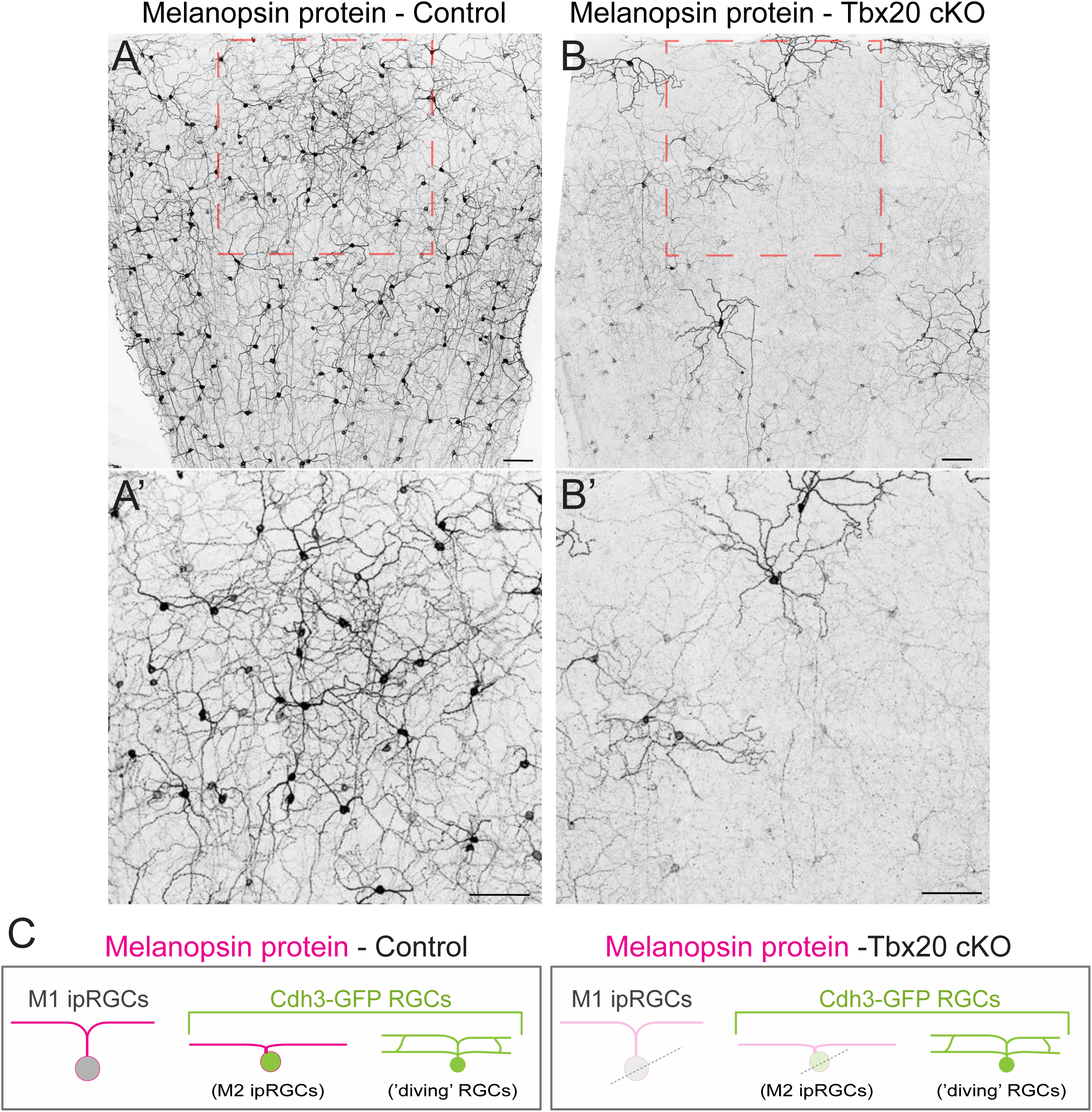
The mosaic pattern of intrinsically photosensitive RGCs that express high levels of melanopsin protein is disrupted in Tbx20 conditional mutants. (A-B’) Photomicrographs showing melanopsin labeling in control (A, A’) and Tbx20 cKO (B, B’) retinas. (C) Schematics depicting reduced melanopsin-expressing RGCs (M1 ipRGCs) in control and Tbx20 cKO retinas. Scale bars: 100μm (A-B’).

To more directly explore the consequence of Tbx20 loss on the organization of the OPN-shell projections we utilized a transgenic mouse line in which a membrane-targeted reporter (Tau-LacZ) was knocked into the Opn4 (melanopsin) locus (Opn4^TauLacZ;^ Hattar et al., 2002, 2006). This reporter line allows for the visualization of M1 ipRGCs and their central projections, including those to the OPN shell. We generated Tbx20 cKO mice on an Opn4^TauLacZ^ background (Cre^+^: Opn4^+/TauLacZ^: Tbx20^f/f^) using a similar breeding scheme as described above. The significantly lower density of β-gal staining in the OPN of these mice also suggests a reduction in the innervation of the OPN (p=0.02; Fig. 5A-H, 5O, schematized in Fig. 5Q). Also, the organization of the M1 ipRGC inputs to the OPN shell was more diffuse in comparison to littermate-control mice (Fig. 5A-H). Furthermore, we found that M1 ipRGC innervation of the SCN, the other major target of these RGCs, was significantly reduced (p<0.001; Fig. 5I-N, 5P). Thus, Tbx20 cKOs have both a loss of Cdh3-RGC inputs to the OPN-core and partial loss of M1 ipRGC inputs to the OPN-shell.

**Figure 5:**
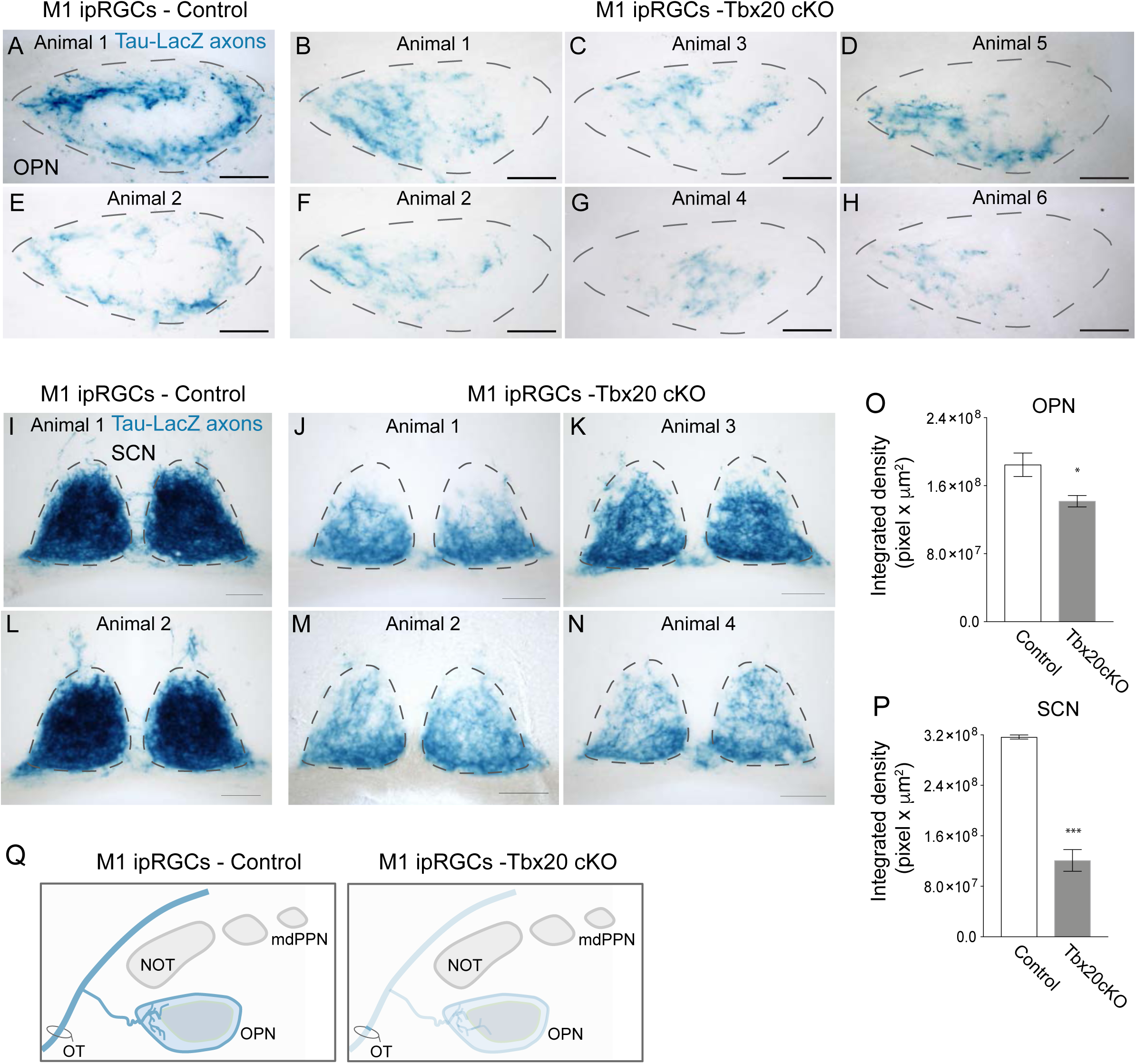
Loss of Tbx20 results in reduced M1 ipRGCs axonal innervation of the OPN and the SCN. (A-H) Photomicrographs showing innervation of M1 ipRGCs (visualized by β-gal staining, blue) in OPN of control (A, E) and Tbx20 cKO mice (B-D, F-H). (I-N) Photomicrographs showing innervation of M1 ipRGCs (visualized by β-gal staining, blue) in SCN of control (I, L) and Tbx20 cKO mice (J, K, M, L). (O) Quantification of density of M1 ipRGC innervation in OPN (β-gal staining density Control Mean ± SEM: 1.85×10^8^ ± 1.38 x10^7^ pixel x µm^2^, n=6 mice; Tbx20 cKO Mean ± SEM: 1.42 ×10^8^ ± 6.74 ×10^6^ pixel x µm^2^, n=6 mice). (P) Quantification of density of M1 ipRGC innervation in SCN (β-gal staining density Mean ± SEM: Control: 3.17×10^8^ ± 3.19 ×10^6^ pixel x µm^2^, n=6 mice; Tbx20 cKO: 1.21×10^8^± 1.72×10^7^ pixel x µm^2^, n=6 mice). (Q) Schematics depicting typical outcomes of M1 ipRGCs innervation of the OPN in control and Tbx20 cKO mice. OPN: olivary pretectal nucleus; SCN: suprachiasmatic nucleus. * p<0.05, ***p<0.001, Student’s t-test. Scale bars: 100μm (A-N).

### Loss of Tbx20 function does not alter the development of other parallel pathways

The Tph2-Cre mouse line (Cre^+^) is a highly useful tool for genetically manipulating Tbx20-RGCs. However, a small fraction of other RGC subtypes (non-Tbx20 expressing) also express Cre recombinase in this mouse line: <3% of image-stabilizing RGCs [Hoxd10-RGCs (Dhande et al., 2013); Seabrook et al., 2017], ∼65% of Cart-expressing RGCs (includes On-Off direction-selective RGCs, Fig. S4A) and ∼37% of alpha-like RGCs (Fig. S4A) express Cre recombinase. Does deletion of Tbx20 impact the development of these optic pathways? We examined the axonal projections of Hoxd10-RGCs in the brains of juvenile (P26) Tbx20 cKO mice (Tph2-Cre^+^; Hoxd10-GFP^+^; Tbx20^f/f^) and littermate-control mice (Tph2-Cre^+^; Hoxd10-GFP^+^; Tbx20^f/+^, and Hoxd10-GFP^+^; Tbx20^f/f^, and Hoxd10-GFP^+^; Tbx20^f/+^) and found that, in the absence of Tbx20 function, Hoxd10-RGCs still innervate their correct targets (Fig. S3A-F). Furthermore, the density of target innervation by the axons of Hoxd10-RGCs mice is not significantly different between Tbx20 cKO mice and control mice (NOT: p=0.64; MTN: p=0.56; DTN: p=0.34, Fig. S3G-I). Next, we stained Tbx20 cKO and control retinas for RGC subtype markers, Cart and SMI32 and found that loss of Tbx20 does not alter the density of either RGC subtype-specific marker (Fig. S4B-G). These data suggest that Tbx20 is required specifically for the development of the non-image-forming visual pathways.

### Loss of Tbx20-RGCs minimally impacts PLR

What is the contribution of Tbx20-RGCs to the PLR? The consensual PLR is defined as the constriction of one eye’s pupil caused by light-stimulating the opposite eye; an effect thought to be mediated by bilateral projections from the OPN to downstream brainstem circuits (Young and Lund, 1994). The ipRGC inputs to the OPN are essential for driving this behavior (Güler et al., 2008; Hatori et al., 2008; Badea et al., 2009; Chen et al., 2011; Sweeney et al., 2014; Mao et al., 2014; Seabrook et al., 2017).

Given the retino-OPN pathway is disrupted in Tbx20 cKO mice, we wanted to know if and how Tbx20-RGCs exert a role in the generation and/or modulation of the consensual pupillary response. Dark-adapted mice were manually restrained and the change in the pupil size of one eye was continuously recorded in response to stimulation of the other eye with a brief (30s) pulse of light. The relative pupil area and percent constriction/dilation were quantified from images extracted from the recording at 5s intervals (see Methods for details). The consensual PLR in Tbx20 cKO mice and littermate controls was tested using a blue LED (470 nm), a wavelength that efficiently stimulates ipRGCs and drives robust PLR responses (Lucas et al., 2001, 2003; Berson et al., 2002; Gamlin et al., 2007). In addition, because the PLR functions over a range of luminance levels, we analyzed consensual PLR at multiple light levels (100 lux, 1,000 lux and 10,000 lux).

First, we computed the average pupil constriction (%) at these three light levels and found that, at 100 lux, the PLR was significantly reduced but the effect was one of only mild attenuation compared to littermate controls (∼ 7% less constricted, p=0.045; Fig. 6A’, 6A”). The same general effect of a significant (but still mild) reduction compared to controls was also observed at 1,000 lux (∼10% less constricted, p=0.0014; Fig. 6B’, 6B”). Pupil constriction was not significantly different between mice lacking Tbx20-RGCs and control mice at 10,000 lux (∼3% less constricted, p=0.06; Fig. 6C’, 6C”). These data indicate that Tbx20-RGCs are mostly dispensable for generating a PLR response. Tbx20-RGCs appear to influence PLR to a small extent only at relatively low light levels (100 lux and 1,000 lux).

**Figure 6:**
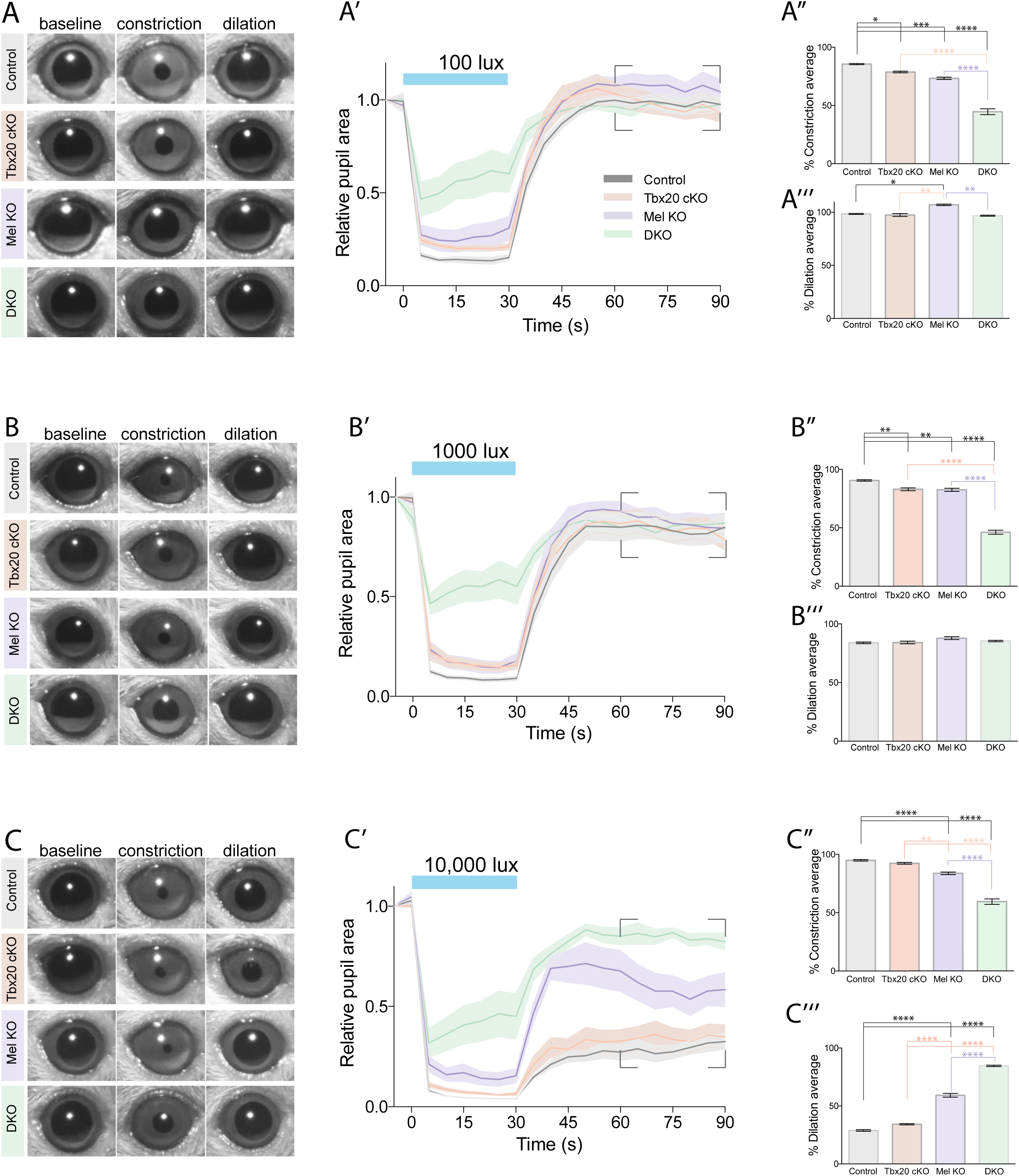
Loss of Tbx20-RGCs minimally impacts the pupillary light reflex. (A, B, C) Representative images of control, Tbx20 cKO, Melanopsin (Mel) KO and Tbx20/Melanopsin double KO (dKO) mice tested with a 100 lux (A), 1,000 lux (B) and 10,000 lux (C) light stimulus at different stages of PLR: baseline (0 s), constriction (30 s), and dilation (60 s). (A’, B’, C’) PLR traces at 100 lux (A’), 1,000 lux (B’) and 10,000 lux (C’). Mean ± SEM (shaded). (A”) Average % pupil constriction in control, Tbx20 cKO, Mel KO, and dKO mice during 100 lux (Control: Mean ± SEM: 85.57 ± 0.45%; Tbx20 cKO: Mean ± SEM: 78.72 ± 0.70%; Mel KO: Mean ± SEM: 73.32 ± 1.04%; dKO: Mean ± SEM: 44.61 ± 2.48%). (B”) Average % pupil constriction in control, Tbx20 cKO, Mel KO, and dKO mice during 1,000 lux (Control: Mean ± SEM: 90.55 ± 0.62%; Tbx20 cKO: Mean ± SEM: 83.03 ± 1.17%; Mel KO: Mean ± SEM: 82.58 ± 1.34%; dKO: Mean ± SEM: 46.25 ± 1.71%). (C”) Average % pupil constriction in control, Tbx20 cKO, Mel KO, and dKO mice during 10,000 lux (Control: Mean ± SEM: 91.90 ± 0.64%; Tbx20 cKO: Mean ± SEM: 89.06 ± 0.75%; Mel KO: Mean ± SEM: 78.82 ± 1.1%; dKO: Mean ± SEM: 53.65 ± 2.27%). (A”’) Average % pupil dilation in control, Tbx20 cKO, Mel KO and dKO mice between 30-60s post-illumination boxed region in A’ at 100 lux (A”’) (Control: Mean ± SEM: 98.38 ± 0.46%; Tbx20 cKO: Mean ± SEM: 97.38 ± 1.39%; Mel KO: Mean ± SEM: 106.9 ± 0.68%; dKO: Mean ± SEM: 96.71 ± 0.48%). (B”’) Average % pupil dilation in control, Tbx20 cKO, Mel KO and dKO mice between 30-60s post-illumination boxed region in B’ in 1,000 lux (B”’) (Control: Mean ± SEM: 83.85 ± 0.69%; Tbx20 cKO: Mean ± SEM: 84.16 ± 1.15%; Mel KO: Mean ± SEM: 87.81 ± 1.19%; dKO: Mean ± SEM: 85.34 ± 0.59%). (C”’) Average % pupil dilation in control, Tbx20 cKO, Mel KO and dKO mice between 30-60s post-illumination boxed region in C’ in 10,000 lux (C”’) (Control: Mean ± SEM: 28.83 ± 0.71%; Tbx20 cKO: Mean ± SEM: 34.21 ± 0.51%; Mel KO: Mean ± SEM: 59.22 ± 1.68%, dKO: Mean ± SEM: 84.59 ± 0.62%). Control: n = 9 mice; Tbx20 cKO: n = 10 mice; Mel KO: n = 7 mice; dKO: n = 6 mice for all conditions. * p <0.05, ** p<0.01, *** p<0.001, **** p<0.0001. Two-way ANOVA with Bonferroni post-hoc test.

OPN-shell projecting M1 ipRGCs are required for generating light-induced pupil constriction (Güler et al, 2008). Given the small but significant decrease in PLR response in Tbx20 cKO mice we wondered whether the near normal PLR response in Tbx20 cKO was driven by the remaining OPN-shell projecting M1 ipRGCs in Tbx20 cKO mice (see above). To dampen the drive of the residual OPN-shell projecting M1 ipRGCs, we generated Tbx20 cKO mice that also lacked the photopigment melanopsin (Mel KO: Opn4 ^TauLacZ/TauLacZ^; see Methods) - hereafter referred to as Tbx20/Mel dKO mice (Tph2-Cre^+^; Tbx20^f/f^; Opn4 ^TauLacZ/TauLacZ^, dKO: double knockout mice). Previous work showed that eliminating melanopsin-photopigment signaling results in a loss of intrinsic photosensitivity in M1 ipRGCs, which in turn leads to reduced pupil constriction in response to light increments (Lucas et al., 2003, Hattar et al., 2003; Panda et al., 2003; Jagannath et al., 2015; Mure et al., 2016). We then tested the consensual PLR in Tbx20/Mel dKO mice. At all three light levels: 100 lux, 1,000 lux and 10,000 lux, pupillary constriction in Tbx20/Mel dKO mice was dramatically reduced relative to mice lacking Tbx20 alone (∼34%, ∼37%, and ∼32%, p<0.0001; less constriction respectively; Fig. 6A’, 6B”, 6B’, 6B”, 6C’and 6C”). Similarly, the PLR of Tbx20/Mel dKO mice was drastically reduced relative to control mice at all three light levels (100 lux: ∼ 41%, 1,000 lux: ∼45%, 10,000 lux: ∼38% less constricted, p<0.0001; Fig. 6A’, 6A”, 6B’, 6B”, 6C’, 6C”).

As discussed above, the PLR is disrupted in mice lacking melanopsin signaling alone. Therefore, we examined whether the severe attenuation in PLR in Tbx20/Mel dKO mice predominantly reflects the loss of melanopsin signaling. The average pupillary constriction (%) in Mel KO mice was significantly less than control mice at 100 lux (∼13% less constricted, p<0.001; Fig. 6A’, 6A”). Similarly, the PLR was significantly attenuated compared to controls at 1,000 lux and 10,000 lux (∼8% less constricted, p=0.0023; and ∼13% less constricted, p<0.0001; respectively; Fig. 6B’, 6B”, 6C, 6C”). Thus, Mel KOs display a significant, although minor, attenuation in PLR at all light levels, a trend similar to that observed in Tbx20 cKOs. Taken together these data indicate that, in Tbx20 cKOs, visual drive provided by the remaining M1 ipRGCs is sufficient for generating PLR. Furthermore, the severity of the PLR loss in Tbx20/Mel double KO mice implies that Tbx20-RGCs cooperatively acts with the OPN-shell pathway to drive the PLR.

Finally, we tested whether the PLR deficits in the mutant mice were somehow a result of abnormal function of the ciliary muscles used for constricting the pupil. Application of the carbachol, a muscarinic agonist that activates the ciliary muscles (Maayani et al., 1975), caused complete pupillary constriction in both control and mutant animals, indicating that ciliary muscle function is unaffected in mutant mice (data not shown).

### Melanopsin- and photoreceptor-mediated-signaling cooperatively act to generate the post-illumination PLR

In addition to studying the pupil constriction that occurs after the onset of a light stimulus, we also systematically analyzed the post-illumination pupillary response (PIPR). The PIPR is the *sustained* pupil constriction after the offset (>30s) of a high intensity, blue light stimulus (Gamlin et al., 2007). In humans and macaque monkeys, the PIPR is primarily mediated by melanopsin signaling (Gamlin et al., 2007; Mure et al., 2009). Sustained depolarization of ipRGCs after the offset of light, due to the slow deactivation of melanopsin, is thought to generate the PIPR (Gamlin et al., 2007). Indeed, a recent report showed that in the presence of a truncated version of melanopsin (lacking residues beyond amino acid 397) that deactivates faster than normal melanopsin, mice fail to maintain pupillary constriction after light offset (Mure et al., 2016).

Following transient illumination of the eye with relatively low-intensity light (100 lux or 1,000 lux) the pupil constricts and then dilates to return to baseline levels (i.e., same as dark adapted levels). This was observed for all the genotypes tested (Controls, Tbx20 cKO, Mel KO, and Tbx20/Mel dKO) (Fig. 6A-6B’’’). However, after transient exposure of the eye to light of a relatively high luminance (10,000 lux) the PIPR occurs at a stage 30-60s after light offset. Similar to the PLR, the disruption in PIPR in Tbx20 cKO mice was modest and not significantly different from controls (∼5% more dilated, p=0.44, Fig. 6C’’’). If melanopsin-signaling alone underlies PIPR, then we would expect Mel KO mice to reach baseline levels of pupil dilation after the offset of an intense light stimulus. Surprisingly, after intense light offset, although the pupils of Mel KOs did dilate significantly more than the pupils of Tbx20 cKOs (∼25% more dilated, p<0.0001; Fig. 6C’’’) the dilation of Mel KO pupils did not reach baseline levels (i.e., complete dilation). These data suggest that, in addition to melanopsin-signaling, other mechanisms influence the PIPR. In the absence of melanopsin-signaling the only source of visual information that can drive any aspect of the PLR or PIPR are light signals encoded by conventional rod/cone photoreceptors relayed to the brain via OPN-projecting RGCs. Indeed, when we eliminated melanopsin signaling and genetically ablated Tbx20-RGCs (Tbx20/Mel dKOs), we found that, on average, the pupils dilate to near baseline levels (∼85% dilation) between 30-60s after the offset of the light stimulus (∼51% more dilated than Tbx20 cKO, p<0.0001; Fig. 6C’’’). Together, these data suggest that not only are Tbx20-RGCs important for the PIPR, but that conventional photoreceptor responses play a significant role in generating the PIPR.

### Targeting Cdh3-RGC using the Cre-DOG system

The loss of both Tbx20-RGC projections to the OPN-core and to the OPN-shell in Tbx20 cKO mice confounds the dissociation of the role of OPN-core versus OPN-shell projecting RGCs in generating PLR. Therefore, to more directly test a role of OPN-core projecting Cdh3-RGCs in the PLR we used a chemogenetic tool hM3Dq [Gq–coupled human muscarinic M3 (hM3Dq) designer receptor exclusively activated by designer drugs (DREADDs)]. hM3Dq is an excitatory DREADD that dramatically increases neuronal firing when bound by its synthetic ligand Clozapine N-oxide (CNO) (Urban and Roth, 2015, schematized in Fig. S5A). Previously, our lab demonstrated that hM3Dq-expressing RGCs increased their spontaneous and light-evoked firing when exposed to CNO (Lim et al., 2016).

To modulate the activity of Cdh3-RGCs, we took advantage of the ‘floxed-Stop-hM3D’ transgenic mouse line that expresses hM3Dq in a Cre-dependent manner (Zhu et al., 2016). To test the efficiency of chemogenetic activation of RGCs in this mouse line, we injected adeno-associated viruses (AAVs) encoding Cre recombinase into the eyes of floxed-Stop-hM3D (Cre^+^/hM3D^+^) mice to achieve widespread expression of hM3D in a random population of RGCs (Fig. S5B, C; n=2 mice). In the absence of CNO, the dark-adapted pupils of Cre^+^/hM3D^+^ mice were completely dilated (Fig. S5D). Upon exposure to CNO, however, the pupils of Cre^+^/hM3D^+^ mice drastically constricted (Fig. S5D). These data show that in floxed-Stop-hM3D mouse the robust activation of RGCs (as measured by the dramatic change in pupil size) occurs only in the presence of both Cre activity and CNO. Our data are in alignment with recent studies demonstrating that activation of hM3Dq expressing RGCs (virally transduced) *in vivo* generates a near complete pupillary constriction (Milosavljevic et al., 2016; Keenan et al., 2016; 2017; Seabrook et al., 2017).

### Selective activation of Cdh3-RGC increases pupil size

Next, to selectively express hM3D in Cdh3-RGCs, we used the Cre-DOG (dependent on GFP) system (Tang et al., 2015) to induce Cre recombinase activity specifically in Cdh3-RGCs (GFP^+^). Cre-DOG is a split Cre system that uses GFP as a scaffold to assemble and induce Cre recombinase activity (schematized in Fig. 7A; Tang et al., 2015). In the retina, AAVs encoding the GFP-dependent split-Cre components (N-Cre and C-Cre) can be used to turn on Cre-dependent-reporter tdTomato specifically in GFP expressing RGCs (schematized in Fig. 7B, 7C). We previously showed that the retrofitting of Cdh3-RGCs using this system is highly selective (Tang et al., 2015). Indeed, we found that in the Cdh3-GFP mouse line nearly all tdTomato expressing RGCs were also GFP^+^ [Fig. 7D-D”; previously quantified: ∼85% of Tdtomato^+^ RGCs are also GFP^+^ (Tang et al., 2015)]. TdTomato-expressing RGC axons in retrofitted Cdh3-GFP mice project to the same targets as Cdh3-RGCs and completely overlap with the axonal projections of Cdh3-RGCs (Fig. 7E-F2”). Importantly, in the absence of GFP (GFP^-^ retinas), no TdTomato expression was observed (previously reported in Tang et al., 2015). Together, these data further validate the high degree of specificity of the Cre-DOG system.

**Figure 7:**
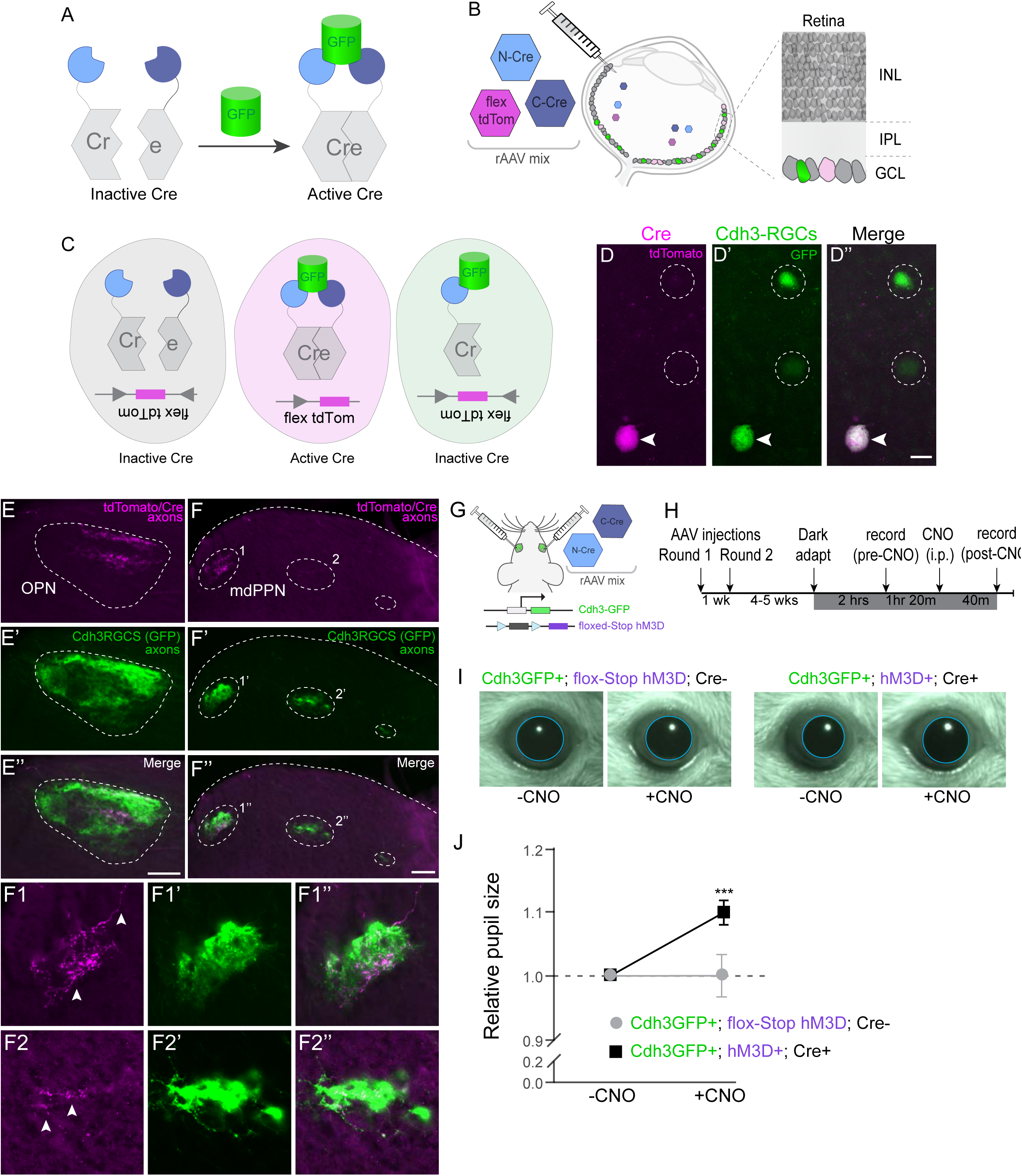
Activation of Cdh3-RGCs increases pupil dilation. (A) Schematic of Cre-DOG (dependent on GFP) system. Complementary split components of Cre bind to GFP thereby inducing Cre recombinase mediate activity. (B) Schematic of ocular injection of viruses encoding the Cre-DOG system (N-Cre and C-Cre) and a Cre-activity reporter (flex-TdTomato). Close up schematic shows resultant tdTomato^+^ RGCs (pink, Cre active), GFP^+^ (green) and GFP^-^ (grey) RGCs. (C) TdTomato reporter is expressed only when a GFP^+^ RGCs are infected by and expresses all three viral components: N-Cre, C-Cre and flex-Tdomato (pink). GFP^+^ neurons (green) that do not express both split-Cre components do express the tdTomato. GFP^-^ neurons (grey) cannot combine the split-Cre system to induce Cre activity. (D-D”) Photomicrographs showing Cdh3-RGC-specific expression of Cre recombinase (visualized via tdTomato reporter). Arrowheads indicate tdTomato^+^ RGCs that are also Cdh3-RGCs (GFP^+^). Dashed circles represent Cdh3-RGCs (GFP^+^) that do not express tdTomato. (E-F2”) Photomicrographs showing the overlap between the projection pattern of tdTomato^+^ RGCs (Cre-DOG targeted) and Cdh3-RGCs. TdTomato^+^ RGC axons terminate in Cdh3-RGC brain targets, the OPN and mdPPN. Dashed white line marks the boundaries of retinorecipient targets. Arrowheads in close up view (F1, F2) indicate tdTomato^+^ axons innervating the mdPPN. (G) Targeting of Cdh3-RGCs using Cre-DOG system. AAV viruses encoding split Cre components (N-Cre and C-Cre) injected binocularly into mice carrying both the Cdh3-GFP and floxed-Stop hM3D transgenes. (H) Experimental timeline. (I, J) Examples (I) and quantification (J) of pupil size before (-CNO) and after (+CNO) administration in Cdh3-RGC targeted (Cdh3GFP^+^; hM3D^+^; Cre^+^; relative pupil size post-CNO Mean ± SEM: 1 ± 0.03; n=9 mice) and control mice (Cdh3GFP^+^; floxed-Stop hM3D; Cre^-^; relative-pupil-size post-CNO Mean ± SEM: 1.1 ± 0.02). See also Figure S5. tdTom: tdTomato reporter; GCL: ganglion cell layer; IPL: inner plexiform layer; INL: inner nuclear layer; OPN: olivary pretectal nucleus; mdPPN: medial division of the posterior pretectal nucleus; hM3D: human M3 muscarinic designer receptor; CNO: Clozapine N-oxide *** p<0.001, paired Student’s t-test. Scale bars: 10μm (D”), 100μm (E”, F”)

To test the role of Cdh3-RGCs in PLR behavior, we bred the Cdh3-GFP mouse line to floxed-Stop-hM3D mouse line to create Cdh3-GFP^+^; floxed-Stop-hM3D mice. Next, we injected a mixture of two AAVs that each individually encode N-Cre and C-Cre (the GFP-dependent split-Cre components) into the eyes of Cdh3-GFP^+^; floxed-Stop-hM3D mice (schematized in Fig. 7G). After 5-6 weeks, the pupils of these mice were recorded prior to, and after, CNO administration (pre- and post-CNO respectively; Fig. 7H). CNO administration alone did not significantly change the pupil size of control animals [Cdh3-GFP^+^; floxed-Stop-hM3D (AAV un-injected); p=0.99; n=9 mice; Fig. 7I, 7J] but CNO mediated activation of hM3D expressing Cdh3-RGCs resulted in a significant increase in pupil size [∼10% increase in Cdh3-GFP^+^; hM3D^+^; Cre^+^ mice; p<0.0001; Fig. 7I, 7J]. Although the activation of Cdh3-RGCs resulted in a highly significant dilation of the pupils the effect is rather small (∼10%). We attribute this minor effect to the low efficiency of the Cre-DOG technique with only a very small fraction of Cdh3-RGCs being converted into Cre-expressing cells (Tang et al., 2015). Regardless, these results suggest that OPN core-projecting Cdh3-RGCs play a unique role in controlling pupil size. Our data suggest a putative “push-pull” system for controlling pupil aperture. The M1-ipRGCs projecting to the OPN-shell drive pupil constriction (push) (Güler et al., 2008 and see above) whereas the Cdh3-RGCs projecting to the OPN-core driving pupil dilation (pull) and thereby dynamically control the full range of PLR behavior. These data, therefore, represent a step towards dissecting the specific function of different types of ipRGCs and in doing so, provide new insight into the distinct functional roles of the different OPN anatomical compartments and pathways.

### Discussion

Our study reveals a previously unrecognized topographically-non-uniform set of RGCs controlling a discrete non-image-forming visual function: pupil size. We also discovered the molecular underpinnings of this circuitry. That, in turn, unveiled mechanistic aspects of their specification and a novel functional component of the visual circuit responsible for pupil reflexes.

### Tbx20 plays a role in establishing non-image-forming parallel optic pathways

Tbx20 is a highly conserved gene both in terms of its structure and function. Tbx20 is known to play an essential role in cardiogenesis in both vertebrate and invertebrate species (Pocock et al., 2008; Jensen et al., 2013; Greulich et al., 2011) and can function as both a transcriptional activator or a transcriptional repressor. Its activity is involved in multiple aspects of cardiac cell proliferation and specification, and structural formation of the heart and mutations in Tbx20 are associated with human congenital heart defects (Cai et al., 2005, 2013; Stennard et al., 2005; Takeuchi et al., 2005; Singh et al., 2005; Sakabe et al., 2012; Kirk et al., 2007; Liu et al., 2008; Posch et al., 2010). The role of Tbx20 in the mammalian nervous system is barely understood, as very limited work has focused on it. Tbx20 was shown by Pfaff and co-workers to be critical for normal development of the hindbrain (Song et al., 2006) by controlling proper migration of facial neurons to their correct destinations in the developing hindbrain, but not required for their initial specification or patterning. Recently, Mason and colleagues identified Tbx20 as a marker of contralateral-projecting RGCs during embryonic development (Wang et al., 2016). Consistent with the findings in this study (Fig. 1 and Fig. S1), their data also shows that Tbx20 is expressed in a subpopulation of RGCs and is graded along the dorsal-ventral axis.

Our results identify Tbx20 as being important for the assembly of non-image-forming retinofugal pathways. In Tph2-Cre mice, Cre recombinase is expressed at ∼E14.5 (see Fig. 2M-R in Seabrook et al., 2017) and Cdh3-RGCs are already born by this developmental stage (Osterhout et al., 2014). Therefore, the loss of Tbx20-RGCs together with the reduction in retinal innervation of non-imaging-forming brain targets in the Tbx20 conditional knockout mice supports the idea that Tbx20 promotes survival rather than specification of different RGC types. Moreover, these differences in phenotypic traits, i.e. loss of neurons in the visual system and migration deficits but not cell loss in the somatosensory system, suggest that Tbx20 can play different roles within different sensory systems. More broadly, this underscores the importance of studying the same genetic program in different sensory systems, as this can reveal new facets and rules governing nervous system development and function.

Previous studies identified the transcription factor T-box family protein Tbr2 (also called ‘Eomes’) and the POU-domain protein Brn3b as necessary for establishing the central projection pathways of non-image-forming RGCs, including all ipRGCs (Quina et al.2005; Badea et al., 2009; Sweeney et al., 2014; Mao et al., 2014; Seabrook et al., 2017). Tbx20 most likely functions downstream of both Tbr2 and Brn3b because i) majority of Tbx20-RGCs express both these factors and ii) Tbx20-RGCs represent a subset of all ipRGCs. Indeed, a recent study by Badea and colleagues (Sajgo et al., 2017) also found that normally Tbx20 is highly enriched in RGCs that express Brn3b but its expression is nearly absent when Brn3b function is lost. In the future, we plan to explore the genes regulated by Tbx20 transcriptional activity as well as the consequences of deleting Tbx20 from the adult visual system.

### Tbx20-RGCs are minor contributors to the constriction phase of the PLR

M1 ipRGCs and their connections to neurons in the OPN shell are critical for driving the pupillary reflex (Güler et al., 2008). Previous studies demonstrated that shell neurons are highly activated in response to light stimuli and that the Edinger Westfal nucleus-which is the downstream target structure of the OPN, receives inputs predominantly from the OPN neurons whose somas reside in the shell (Prichard et al., 2002; Baver et al., 2008). Although ablating all shell-projecting M1 ipRGCs leads to a severe attenuation of the PLR (∼42% reduction in pupil constriction at high light levels, and no constriction at low light levels), light-induced pupil constriction is not completely abolished in the absence of M1s (Güler et al., 2008). Complete loss of the PLR only occurs when all ipRGCs, as well as rod/cone mediated signaling, are eliminated (Hattar et al., 2003; Panda et al., 2003; Badea et al., 2009; Hatori et al., 2008, Sweeney et al., 2014; Mao et al., 2014). Removing M1s and all rod/cone signaling is effectively the same as removing all RGCs. Thus, RGCs other than M1 ipRGCs projecting to the OPN shell must contribute to the PLR.

Conditionally knocking out Tbx20 eliminated a subset of RGCs that project to the OPN core as well as the OPN shell. Alone this manipulation did not lead to a severe deficit in light-evoked pupillary constriction. This is not surprising, as the population of M1 ipRGCs that do not express Tbx20 remain intact in Tbx20 cKO mice. Indeed, even an extremely small number of retinal inputs to the OPN (∼7-20%) is sufficient to drive a strong PLR response (Güler et al., 2008; Mao et al., 2014; Seabrook et al., 2017). A key contribution of Tbx20-RGCs to the pupillary reflex was revealed, however, when melanopsin-mediated signaling was abolished in Tbx20 cKO mice. Depending on the intensity of the light stimulus, pupillary constriction in those mice was reduced ∼45-55% in Tbx20/Mel dKO mice as compared to control mice. The level of pupillary constriction deficits observed in Tbx20/Mel dKO mice are comparable to those observed when all shell-projecting M1 ipRGCs are ablated (Güler et al., 2008). Our experimental data cannot precisely disambiguate the specific role of OPN-core versus OPN-shell in the PLR, as retinal innervation to both these targets is compromised in Tbx20 cKO mice. However, the most parsimonious explanation for the dramatic disruption of the PLR in Tbx20/Mel double KO is that it is primarily due to the loss of M1 ipRGC (Tbx20^+^) innervation to the OPN-shel,l compounded by the loss of melanopsin signaling in the remaining OPN-shell projecting M1 ipRGCs (Tbx20^-^).

### Conventional-photoreceptor driven visual signals contribute to the PIPR

Our analysis of the pupillary reflex in Tbx20/Mel dKO mice unveiled a novel role of rod/cone mediated signaling in generating the post-illumination pupillary reflex (PIPR)-which is the persistent constriction of the pupil after the offset of a relatively high-intensity light stimulus. The PIPR is widely used as an indicator of ipRGC health in humans and non-human primates as a diagnostic for a broad range of diseases including glaucoma, multiple sclerosis, and seasonal affective disorder (Meltzer et al., 2017, Feigl & Zele 2014). The PIPR is thought to be driven by the sustained activity of ipRGCs after the offset of light. Indeed, a recent report showed that in the presence of a truncated version of melanopsin lacking residues beyond amino acid 397, mice fail to maintain pupillary constriction after light offset (Mure et al., 2016). The circuit-based mechanistic underpinnings of the PIPR has not been examined. We found that similar to primates, the PIPR of mice is strongly dependent on melanopsin signaling, as Mel KO failed to maintain a sustained pupillary constriction after the offset of a relatively high irradiance stimulus. However, rodent ipRGCs show persistent activity, albeit attenuated, even in the absence of the melanopsin signaling (Schmidt et al., 2014; Wong 2012). Indeed, we found that the pupillary escape from constriction after light offset was significantly greater in the Tbx20/Mel dKO mice in comparison to mice lacking only melanopsin signaling. This suggests that OPN-projecting ipRGCs use both melanopsin-mediated as well as rod/cone-mediated visual signals to generate the PIPR.

### Insights into the role of the OPN ‘core’ pathway

The OPN receives inputs from multiple types of RGCs namely, M1 and M2 ipRGCs and from ‘diving’ RGCs. Each of these RGC types is capable of encoding and relaying irradiance changes necessary for driving the PLR. However, our data and the work of others indicate that M1 ipRGC inputs to the shell provide the main drive for pupillary constriction and are sufficient to drive near normal PLR. This raises the question of the purpose of the retina-OPN-core pathway. We selectively probed the functional output of the OPN-core projecting Cdh3-RGCs by reversibly manipulating their activity using chemogenetic tools and found that Cdh3-RGC activation, in fact, results in a modest dilation of the pupil. This suggests that the OPN-shell and OPN-core function in a “push-pull” manner to modulate the aperture of the eye. Indeed, OPN inputs and outputs of extremely heterogeneous across multiple species (Gamlin 2006). The M1 ipRGC-to-OPNshell-to-EW pathway is part of the parasympathetic system and has received a lot of attention due to its role in generating light-induced pupil constriction. However, OPN neurons also project to several other nuclei including the SC, periaqueductal gray, nucleus and nucleus of the posterior commissure, brain targets have previously been implicated as part of the sympathetic pathway involved in controlling pupil dilation and arousal states (Klooster et al., 1995; Wang and Munoz, 2015; Salay et al., 2018). In future work, it will be important to dissect the downstream targets and other inputs to OPN-core neurons as a way to parse the contributions of this circuit to pupil size dynamics in the context of both light-mediated and light-independent arousal.

## Methods

### Animals

Experimental procedures were in accordance with NIH guidelines and approved by the Institutional Animal Care and Use Committees at UCSD and Stanford University School of Medicine. The generation and characterization of Tbx20 floxed and Tbx20-GFP BAC transgenic mouse lines were described previously (Cai et al., 2005 and Shen et al., 2011, respectively). Tryptophan hydroxylase 2-Cre (Tph2-Cre) mice were obtained from MMRRC (https://www.mmrrc.org/catalog/sds.php?mmrrc_id=36634) (Osterhout et al., 2011; 2014). Cadherin-3-EGFP (Cdh3-GFP) were obtained from MMRRC and previously characterized (Osterhout et al., 2011; 2014). Homeobox d10 BAC-EGFP (Hoxd10-GFP) mice were obtained from MMRRC and described previously in Dhande et al., 2013. Opn4 (opsin 4/melanopsin) Tau-LacZ mice were obtained from Jackson Laboratory (https://www.jax.org/strain/021153) and reported previously (Hattar et al., 2002; 2006). Opn4 Tau-LacZ homozygous mice do not express melanopsin (Mel KO). CAG-LSL-hM3Dq/mCitrine transgenic mice were obtained from the Jackson Laboratory (https://www.jax.org/strain/026220) and reported previously (Zhu et al., 2016). These mice express the hM3Dq DREADD and mCitrine reporter after Cre-recombinase mediated excision of a floxed-Stop cassette.

### Histology & Immunohistochemistry

Mice were transcardially perfused, brains harvested and postfixed overnight (∼12hrs) in 4% PFA at 4°C, immersed in 30% sucrose for 2-3 days at 4°C, and sectioned at 45μm on a sliding microtome. Tissue was incubated overnight at 4°C in primary antibodies and incubated for 2hrs at room temperature with appropriate secondary antibodies. Retinas were harvested and processed for immunohistochemistry as previously described (Huberman et al., 2008; Tang et al., 2015).

Primary antibodies used were: rabbit anti-GFP (1:1,000, Molecular Probes Cat# A-6455, RRID:AB_221570), chicken anti-GFP (1:1,000; Aves Labs Cat# GFP-1020, RRID:AB_10000240), rabbit anti-Cart (1:1,000, Phoenix Pharmaceuticals Cat# H-003-62, RRID:AB_2313614), mouse SMI-32 (1:1,000, Covance Research Products Inc Cat# SMI-32P, RRID:AB_2314912), rabbit or guinea-pig anti-Tbx20 (1:1,000; Song et al., 2006), rabbit anti-melanopsin (1:1,000; Advanced Targeting Systems Cat# AB-N39, RRID:AB_1608076), rabbit anti-Tbr2 (1:1,000, Abcam Cat# ab23345, RRID:AB_778267), mouse anti-β-Galactosidase (β-Gal) (1:1,000, Sigma-Aldrich Cat# G8021, RRID:AB_259970), rabbit anti-dsRed (1:1,000, Clontech Laboratories, Inc. Cat# 632496, RRID:AB_10013483), rabbit anti-Brn3b (1:1,000, Quina et al., 2005). Species-specific secondary antibodies conjugated to Alexa Fluor 488, 594 or 647 (1: 1,000; Invitrogen and Jackson Laboratory) were used.

### X-gal staining

Tissue was harvested and sectioned as described above, with the exception that brains were fixed for 4hrs in 4%PFA. X-gal staining was performed as previously described (Dhande et al., 2012). Control and Tbx20 cKO mice that were also heterozygous for the Opn4 Tau-LacZ allele were used for X-gal staining.

### RGC marker and density quantification

Quantification of GFP^+^ RGCs and expression of molecular markers was performed as previously described (Dhande et al., 2013). For marker analysis, labeled RGCs were counted in both the central and peripheral region of a randomly selected retinal leaflet. Counts were performed by manual cell identification using Neurolucida (MicroBrightField).

### In situ hybridization and quantification

Mice were transcardially perfused, retinas harvested and postfixed overnight in 4% PFA at 4°C, immersed in 30% sucrose for one day at 4°C, embedded in Tissue-Tek O.C.T. compound (Sakura) and sectioned at 15μm on a cryostat. RNAscope Multiplex Fluorescent Assay was performed according to manufacturer’s instructions using custom Tbx20 probes (targeting the 460-1708 region of NM_194263.2; Cat No. 511991; Advanced Cell Diagnostics). Images were acquired with a Zeiss LSM 880 or a Zeiss AxioScan microscope. Fluorescently labeled cells were manually counted from 5 consecutive sections per animal.

### Intravitreal injections of tracers or viruses

Binocular injections of 1-2μL of Cholera Toxin-β conjugated to Alexa Fluor488 (CTβ - 488, Invitrogen) or AAV-iCre (Vector BioLabs) or a 1:1 mix of AAV2/1.EF1a.N-CretrcintG (N-Cre, 3.95×10^13^ GC/mL) and AAV2/1.EF1a.C-CreintG (C-Cre, 3.94×10^13^ GC/mL, Gene Transfer Vector Core, Harvard Medical School). CTβ-488 injected mice were killed two days later, and virus injected mice were killed after behavioral analyses. Brain tissue and retinas were harvested and processed as described above (see *Histology & immunohistochemistry*).

### Retina reconstructions and cell counts

Flat mount retinas with labeled Tbx20-GFP or Cdh3-GFP cells were outlined in Image J to demarcate tear locations. Cells were identified based on the signal intensity and cell body positions were recorded using the ImageJ ROI manager. These parameters were imported into the Retistruct package in R (Sterratt et al., 2013; R Development Core Team, 2014). Flat mount retinas were then reconstructed to standard spherical retina space and labeled cells were plotted in spherical coordinates. Isodensity contour lines were derived using kernel density estimates. Cell counts were obtained from each retinal pole. For comparing cell counts in opposing retinal poles, binomial tests with the stats R package were performed for each retina. To compare average cell counts in opposing poles, paired t-tests were performed.

### Quantification of target area and innervation density

Measurements were taken from 3-5 tissue sections depending on the target size. CTβ–488 or GFP labeling was used to outline the boundaries of the retinorecipient targets. A template boundary based on representative control OPN and SCN boundaries was used for measuring β-Gal staining density. ImageJ ‘Area’ and ‘Integrated density’ tools were used to measure the retinorecipient target area and density of innervation (product of mean gray pixel value and target area).

### Pupillometry

The consensual PLR of dark-adapted mice was recorded and quantified as follows. After 2hrs of dark-adaption, under infrared lighting conditions, the left eye of the awake hand-restrained mouse was illuminated with a light intensity of 100, 1,000 or 10,000 lux using a single 470 nm tunable LED (Thor Labs, M470L3) for 30s. The consensual (right eye) was recorded using a Sony handycam immediately before illumination, during illumination (30s), and for 1 minute after the LED turned off. 3-4 recordings for each lighting condition were made from each mouse between Zeitgeber time (ZT)6 and ZT10 (where ZT0 is lights-on and ZT12 is lights-off). To induce complete ciliary constriction, 100 mM Carbachol was prepared and applied as previously described (Lucas et al., 2003).

For quantification of pupil size, individual frames were extracted from the video recordings at every 5s from t= −5 sec (dark adapted, lights off), to t= 90 sec. Pupil diameter was calculated for each of these frames and normalized to the diameter of the eye in the same frame in ImageJ. Pupil area was calculated for each time point relative to the initial dark-adapted pupil size at t= −5 sec. Chemogenetic induced changes in pupil size were recorded and quantified as described above, with the following modifications: under constant dark conditions (no LED stimulus) images of the right pupil were captured before and 40 minutes after intraperitoneal administration of Clozapine N-oxide (CNO, 5mg/kg, Tocris Bioscience). Post-CNO pupil size was normalized to pre-CNO pupil size.

### Targeting Cdh3-RGCs using the Cre-DOG system

The Cre-DOG system was used to selectively infect Cdh3-GFP RGCs with Cre as in: Tang et al. (2015). Briefly, intraocular injections of three AAVs each encoding either N-Cre, C-Cre or flex-TdTomato-reporter, were made into 2-3 month old Cdh3-GFP mice. 2-4 weeks later, retinas and brains were harvested and processed as described above (see *Histology & immunohistochemistry*). For chemogenetic experiments, 1-2μL of a 1:1 combination of AAV N-Cre and AAV C-Cre viruses were injected binocularly into mice carrying both the CAG-floxed-Stop-hM3D/mCitrine transgene and the Cdh3-GFP transgene. Two rounds of retinal injections were performed one week apart. Pupillometry was performed 5-6 weeks after the last set of retinal injections animals.

### Experimental design and statistical analysis

For behavioral analyses, 2-3 month old mice were used. For all other experiments, mice aged postnatal day (P)19-P28 were used, unless otherwise stated. All mice were maintained on a mixed background. Mice of both sexes were used in all experiments. All data are expressed as Mean ± Standard Error of Mean (SEM). Data were significant when p<0.05, as determined by a two-way ANOVA with Bonferroni post-hoc test for PLR analysis, and by Student’s t-test for all other analyses, using GraphPad (Prism program, California, USA). Paired tests were used to compare within-group repeated measure data.

## Author Contributions

O.S.D. and A.D.H conceived and designed the experiments and wrote the paper. O.S.D. and T.A.S. characterized the Tph2Cre mouse line. O.S.D., A.H.P., N.I., and P.L.N. performed brain histology. O.S.D., A.H.P. and N.I. performed behavior experiments. O.S.D and A.H.P. imaged and analyzed histology and behavior data. L.D.S. analyzed retinal gradients. J.T.W. performed initial molecular screen. S.E. provided the Tbx20 conditional allele mice and Tbx20-GFP mice. O.S.D. performed retinal histology; performed and analyzed chemogenetic experiments; and prepared the figures and manuscript.

## Acknowledgements

This work was supported by a Knights Templar Eye Foundation Grant (O.S.D.), NIH RO1 EY022157 (A.D.H.), NIH RO1 EY026100 (A.D.H.), and a Pew Scholar Award (A.D.H.). We thank Dr. Samuel Pfaff and Dr. Eric Turner for kindly providing Tbx20 and Brn3b antibodies and Dr. Rana El-Danaf and Dr. Supraja Varadarajan for helpful comments on an earlier draft on the manuscript.

## Declaration of Interests

The authors declare no competing interests.

## Supplemental Information

**Figure S1, related to Figure 1:**
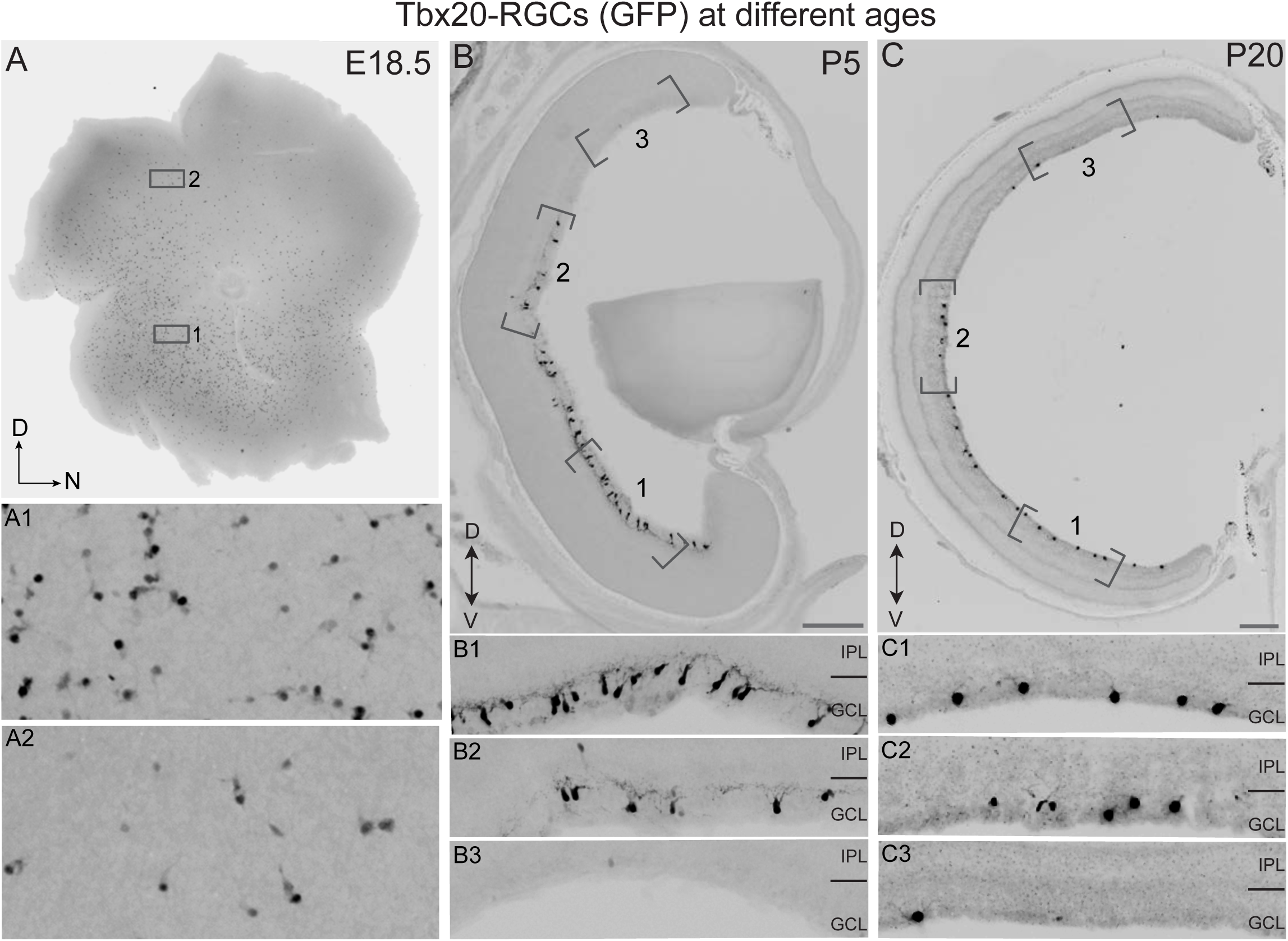
Tbx20 is highly enriched in a group of uniquely distributed non-image-forming RGCs that include Cdh3-RGCs. (A-A2) Flat mount retina showing Tbx20-RGCs (black) are arranged in a ventral/high to dorsal/low gradient of distribution at E18.5. (B-C3) Retinal cross-sections showing ventral/high to dorsal/low gradient of distribution of Tbx20-RGCs (black) at P5 and P20. D: dorsal; N: nasal; V: ventral; GCL: ganglion cell layer; IPL: inner plexiform layer. Scale bar: 200μm (B, C).

**Figure S2, related to Figure 1:**
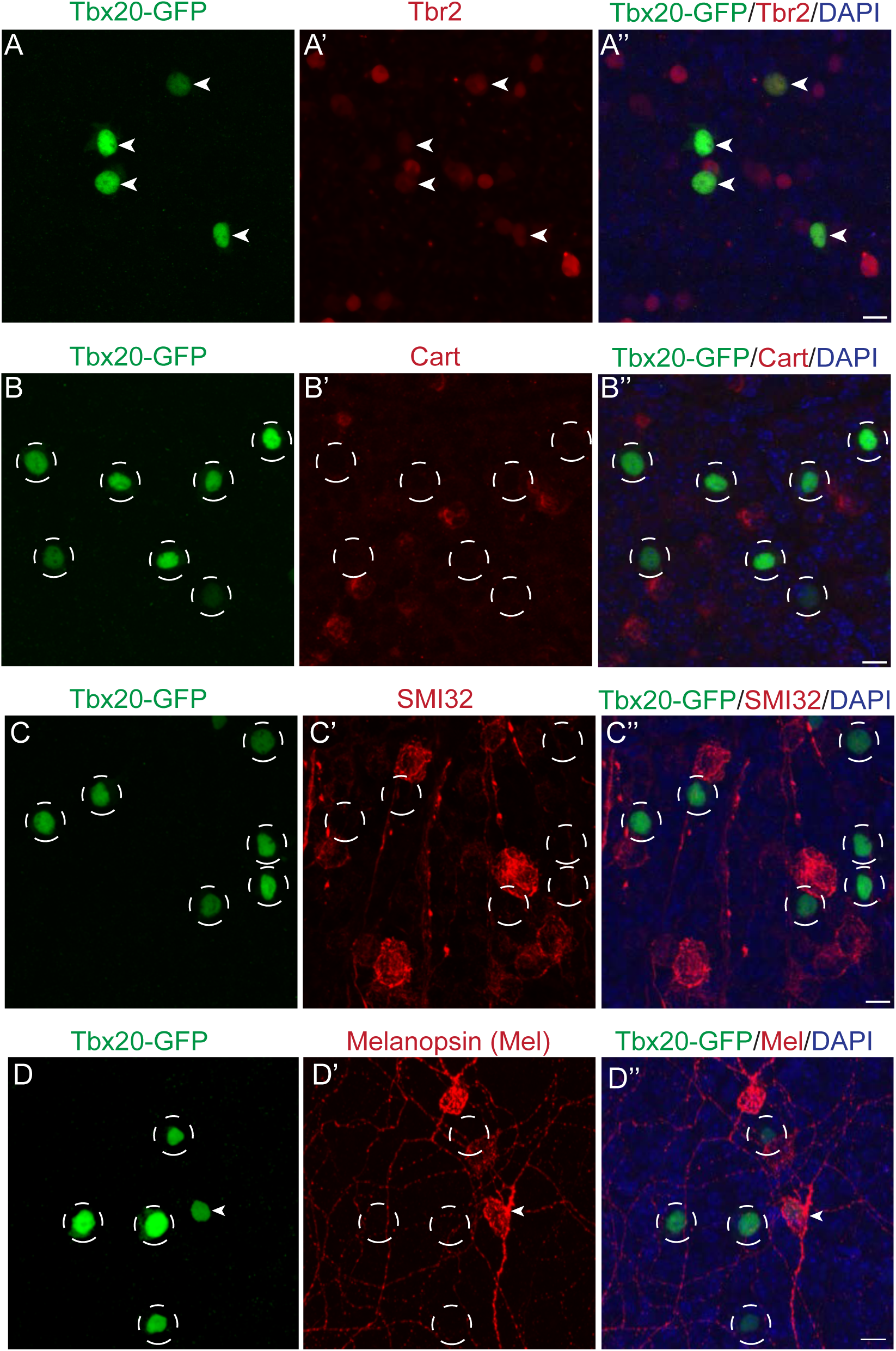
RGC type-specific molecular marker analysis suggests Tbx20-RGCs are part of the non-image-forming RGC family. (A-D”) Photomicrographs showing Tbx20-RGCs (GFP) stained with Tbr2, a marker of non-image-forming RGCs. Arrowhead indicates overlap of GFP and Tbr2 and open circles indicate lack thereof. (B-C”) Photomicrographs showing Tbx20-RGCs (GFP) stained with Cart, a marker for direction-selective RGCs or SMI-32, a marker for alpha-like RGCs. Open circles indicate a lack of overlap between markers. (D-D”) Photomicrographs showing Tbx20-RGCs (GFP) stained with Melanopsin. Arrowhead indicates overlap of GFP and melanopsin. Open circles indicate a lack of overlap between these two markers. Scale bar: 10μm (A”, B”, C”, D”).

**Figure S3, related to Figure 2:**
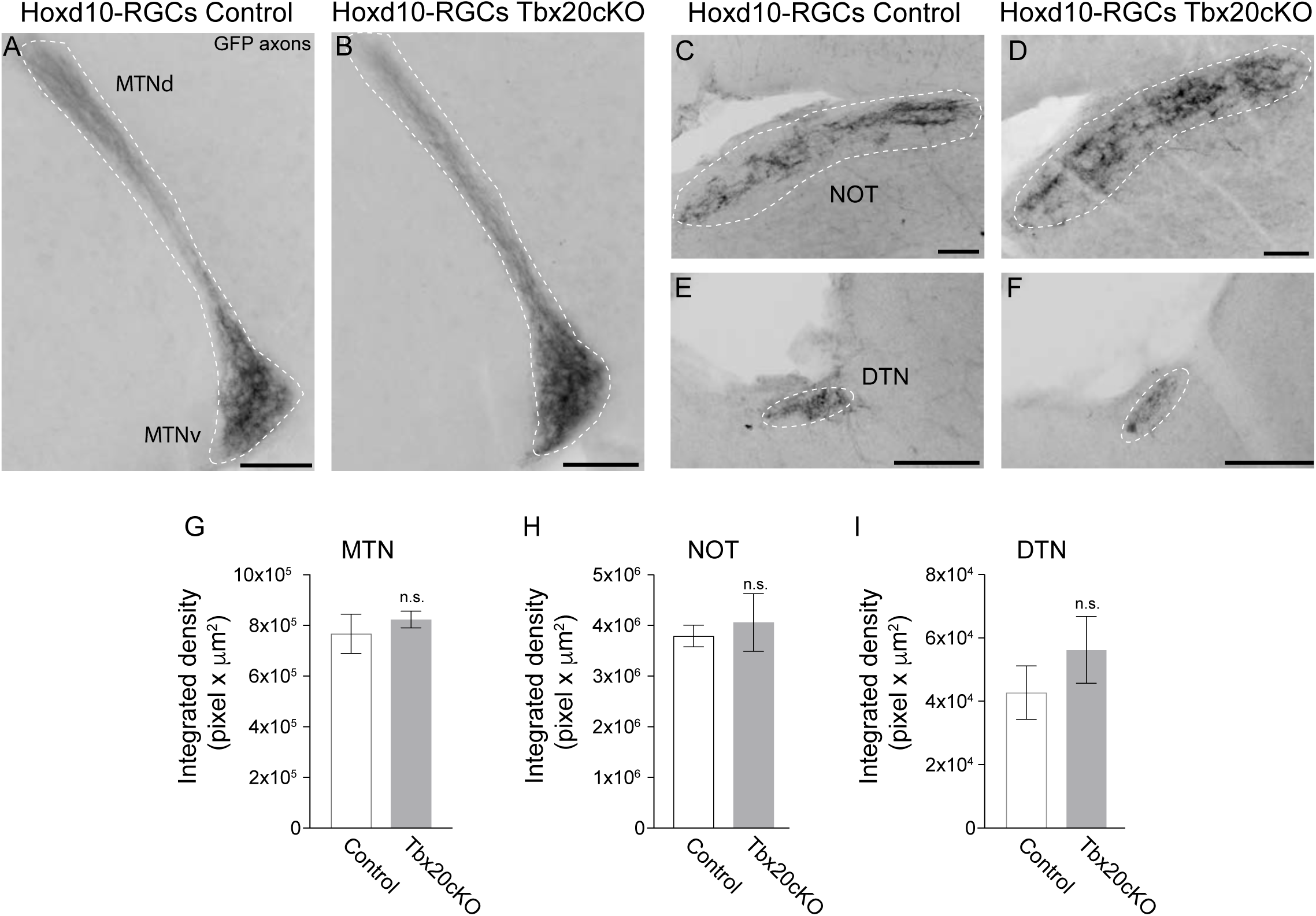
Conditional deletion of Tbx20 from accessory optic system projecting RGCs does not alter their target innervation-density. (A-F) Photomicrographs showing Hoxd10-RGC axonal projections (GFP, black) to the targets comprising the accessory optic system, MTN, NOT and to the DTN, in a littermate control mouse (A, C, E) and Tbx20 cKO mouse (B, D, F). (G-I) Quantification of density of Hoxd10-RGC axonal innervation in MTN (G) (Control Mean ± SEM: 7.7 x10^5^± 7.7×10^4^ pixel x µm^2^; Tbx20 cKO Mean ± SEM: 8.2 ×10^5^± 3.3×10^4^ pixel x µm^2^), NOT (I) (Control Mean ± SEM: 3.8 x10^6^± 2.2 ×10^5^ pixel x µm^2^; Tbx20 cKO Mean ± SEM: 4.1 ×10^6^± 5.7×10^5^ pixel x µm^2^) and DTN (I) (Control Mean ± SEM: 4.3 ×10^4^± 8.4×10^3^ pixel x µm^2^; Tbx20 cKO Mean ± SEM: 5.6 ×10^4^± 1.1×10^4^ pixel x µm^2^) in control (n=5 mice) and Tbx20 cKOs (n=4 mice). MTNd: dorsal medial terminal nucleus; MTNv: ventral MTN; NOT: nucleus of the optic tract; DTN: dorsal terminal nucleus. n.s.: not significant, p>0.05. Student’s t-test. Scale bars: 100μm (A-D), 50μm (E, F).

**Figure S4, related to Figure 2:**
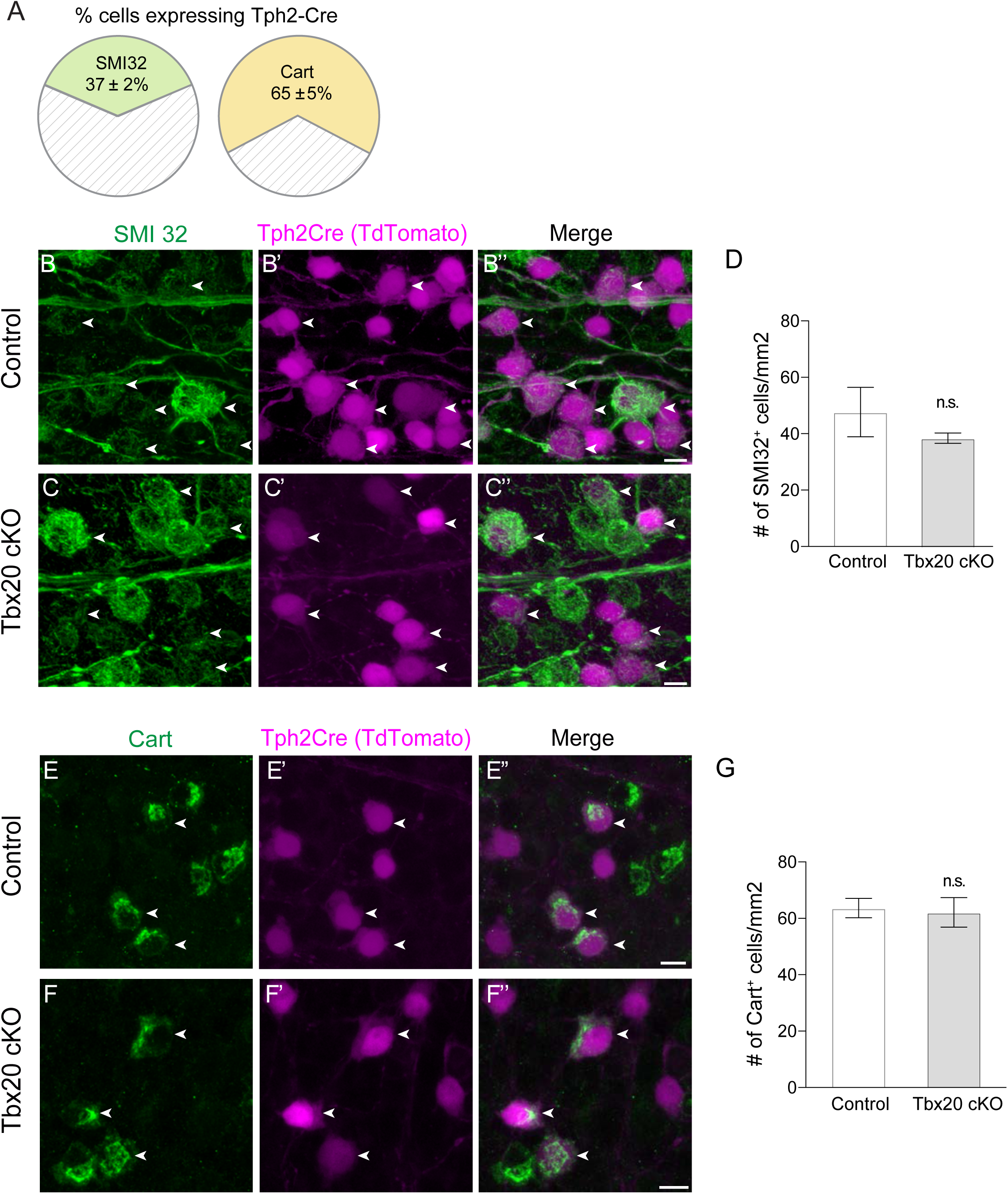
Loss of Tbx20 in other RGC subtypes does not alter their density. (A) Quantification of SMI-32^+^ cells that express Tph2-Cre (n=3 retinas/mice; 144 SMI32^+^ cells) and Cart^+^ cells that express Tph2-Cre (n=5 retinas/mice; 321 Cart^+^ cells) at P25. Data shown are Mean ± SEM. (B-C”) Photomicrographs showing SMI32 and Tph2-Cre (tdTomato) expression in control and Tbx20 cKO mice. White arrowheads indicate co-expression. (D) Quantification of density of SMI32+ RGCs in control and Tbx20 cKO mice (Control Mean ± SEM: 48 ± 9 cells/µm^2^, n=3 mice; Tbx20 cKO Mean ± SEM: 39 ± 2 cells/µm^2^, n=3 mice). (E-F”) Photomicrographs showing Cart and Tph2-Cre (tdTomato) expression in control and Tbx20 cKO mice. White arrowheads indicate co-expression. (G) Quantification of density of Cart+ RGCs in control and Tbx20 cKO mice (Control Mean ± SEM: 64 ± 3 cells/µm^2^, n=5 mice; Tbx20 cKO Mean ± SEM: 63 ± 5 cells/µm^2^, n=3 mice). n.s.: (not significant): p>0.05. Student’s t-test. Scale bar: 10μm (B”, C”, E”, F”)

**Figure S5, related to Figure 7:**
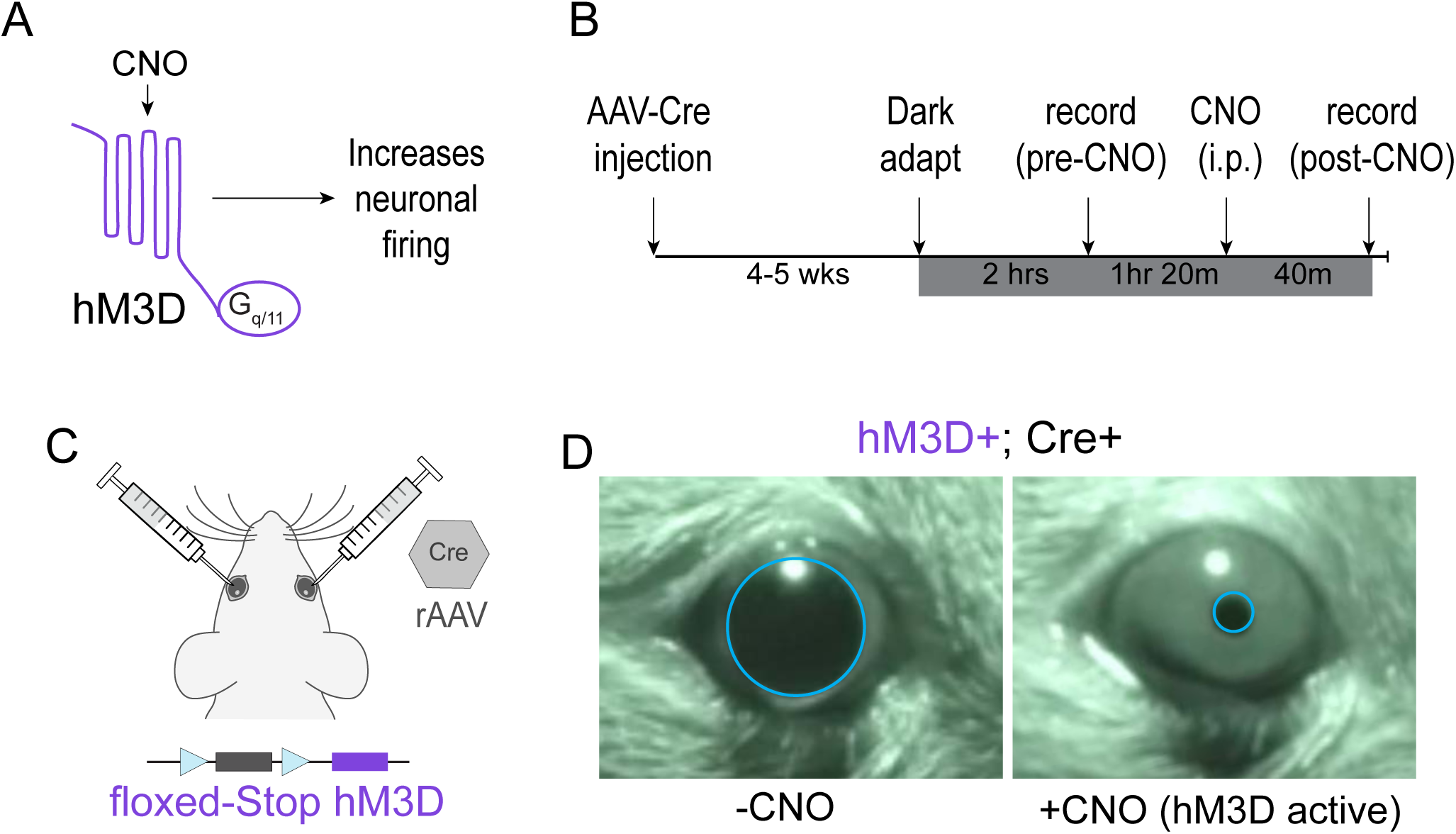
Chemogenetic activation of RGCs generates robust pupil constriction. (A) Schematic of Gq coupled hM3D receptor. (B, C) Experimental timeline (B). AAVs encoding Cre recombinase were injected into both eyes of floxed-STOP hM3D mice (C). 4-5 weeks post-injection pupil size was measured after dark adaptation before and after intraperitoneal (i.p.) CNO (Clozapine N-oxide) administered (B). (D) Example of pupil size pre (left) and post (right) CNO administration.

